# PlantLRR-PRR: A Reproducible Annotation Pipeline Reveals Contrasting Evolution of Developmental and Defense Receptor-Like Proteins in Tomato

**DOI:** 10.64898/2026.09.08.750299

**Authors:** Nandeesh Jalahalli Rangegowda, Remco Stam

## Abstract

Plants perceive external and internal signals via receptors to modulate their growth, development, and defenses. Leucine-rich-repeat receptor-like-kinase (RLKs) and receptor-like-proteins (RLPs) are two major cell-surface receptors in plants. RLPs are a unique gene family (well-studied and characterized) and play an important role in plant development and defense activities against pests and pathogens. Yet their comparative analyses are hampered by a lack of reproducible and incomplete annotation tools. Here we present the PlantLRR-PRR, a reproducible and standardized RLP/RLK annotation pipeline. It outperforms previous tools and helps to dissect the RLP variation among *Solanum spp*.

We investigated RLP diversity in eight genomes from five wild tomato *Solanum* sect Lycopersicum. We found limited intra- but moderate inter-specific copy number variation displaying a possible long-term diversification (gain and loss) of RLPs driven by host, environment, and pathogen interactions. Interestingly, we observed a dual-evolutionary pattern characterized by conservation and diversification of developmental- and defense-related RLPs, respectively. Overlapping with this, we found transposable elements (TEs) highly enriched around defense-related RLPs, supporting a strong role of TEs in promoting loss, gain, and structural variations. Further, zooming into the known Cf5 and CuRe1 RLP gene cluster’s revealed signatures of typical birth–death patterns. Comparative analysis of RLPs in wild tomato species revealed RLP diversity that is not apparent from cultivated tomato alone, highlighting the value of wild germplasm for understanding RLP family evolution. Together, our results provide a comparative framework for understanding the evolutionary divergence and conservation in the RLP family and establish a foundation for linking receptor evolution with functional resistance, and can form a stepping stone for translational applications in crop improvement.

## Introduction

A plant’s ability to detect and respond to environmental cues underlies its adaptation and evolution over time. Plants have evolved with receptors to modulate their growth, development, and defense by recognizing internal and external signals. Lacking adaptive immunity, plants have evolved with an innate immune system primarily consists of cell-surface pattern-recognition receptors (PRRs) and intercellular receptors to perceive apoplastic and cytoplasmic deposited pathogen signals, respectively (Jones & Dangl, 2006; Ngou *et al*., 2022a). Focusing on PRRs, all functionally characterized receptors carry a signal peptide (SP) to guide the protein movement into the apoplastic region, an extracellular domain (ECD) to perceive pathogen signals, and transmembrane domains (TMDs) to attach to the plasma membrane (Wang *et al*., 2008; Macho & Zipfel, 2014; Snoeck *et al*., 2023). There are different types of ECDs having PRRs, and Leucine-rich repeats (LRRs) are the most prominent ones. There are two major types of LRR-PRR families, LRR receptor-like-kinase (RLKs) and receptor-like-proteins (RLPs). Along with SP, LRRs and TMD, RLKs have an additional cytoplasmic kinase domain and RLPs have a short cytoplasmic tail at their C-terminal (Macho & Zipfel, 2014).

Plant receptors can be divided into developmental and defense-related, purely based on functional attributes. Developmental receptors are mostly conserved across plant species (i.e. both on a phylogenetic and molecular scale) (Peterson *et al*., 2010; Krusell *et al*., 2011; Steidele & Stam, 2021). Defence-related are said to be under stronger evolutionary pressures to diversify, mainly from the highly dynamic incoming phytopathogens (Muir *et al*., 2014; Torres Ascurra *et al*., 2023). As pathogens also need to avoid recognition, in the long run this has the potential to turn into a arms race, a well-established concept in plant-microbe interactions, can be interpreted that the plant’s immune system is co-evolving with the disease-causing molecules (effectors) from the phytopathogens (Ngou *et al*., 2022a). Nonetheless, inconsistent pressure from phytopathogens and multiple sources on host plants also creates an opportunity to lose or gain receptors across the plant population and species, creating diversity for different receptor types. This was well-studied for the intercellular nucleotide-binding LRR (NLRs) gene family evolution across multiple studies targeting *Arabidopsis* and *Solanum spp.* (Michelmore & Meyers, 1998; Van De Weyer *et al*., 2019; Stam *et al*., 2019; Sutherland *et al*., 2024; Silva-Arias *et al*., 2025). However, a comparative frame for understanding PRRs, especially RLPs, has been overlooked, as most focus lies on the larger NLRs and RLK families. RLPs might unveil distinct patterns of gene family evolution across plant populations and species, especially because they possess a dual role that can be systematically inferred from their sequence (Steidele & Stam, 2021). Two well-known developmental-related RLPs, Too-many-mouth (TMM) involved in stomatal patterning, and Clavata-2 (CLV2) for the maintenance of meristematic tissue were investigated and found to be highly conserved and present in single copies across plant species (Peterson *et al*., 2010; Krusell *et al*., 2011; Steidele & Stam, 2021). Cf9 an Cf4 are most likely the best studied defence associagted RLPs in tomato, recognizing the *Cladosporium fulvum* (now *Passalora* fulva) effectors Avr9 an 4 resp. and triggering strong hypersensitive response associated defence (Jones *et al*., 1994; Thomas *et al*. 1997). Previous work on *S. chilense* has shown presence-absence variation of potential RLPs against *Cladosporium fulvum* effectors, including Avr9, Avr4, and Avr-mix (a complete set of *Cladosporium fulvum* secreted apoplastic effectors) across different populations. But the totality of RLP diversity among different populations and species of wild and cultivated tomatoes is yet to be conducted (Stam et al., 2019).

Immune-related RLPs sense conserved pathogen-associated molecular patterns (PAMPs) and pathogen effectors in the apoplastic region using LRR domains. Upon perception, they activate pattern-triggered immunity against PAMPs and effector-triggered immunity against effectors (Jones & Dangl, 2006; Ngou *et al*., 2022a). Unlike RLKs, which transmit downstream signalling via their own kinase domain, RLPs solely coordinate with co-receptors like SOBIR-1 (suppressor of BIR-1) to initiate downstream signalling (Van Der Burgh *et al*., 2019; Seifbarghi *et al*., 2020; Yang *et al*., 2020). Well-known defense RLPs are composed of conserved domains named from A to G based on respective motif sequences. A and B are putative SP and cysteine rich domins. C is the ECD, further divided into C1 to C3 subsections, where C1 and C3 are LRR-domains and C2 is the co-called Island domain, consisting of a Yx8KG motif (Snoeck et al., 2023). Domains D, E, and F are spacer, an acidic subdomain, and a TMD, respectively. At last, domain G is a short cytoplasmic tail without a kinase domain (Jones *et al*., 1994; Snoeck *et al*., 2023). A few functionally well-characterised defence-related RLPs include Cf9, Cf5 (for Avr5), Cf4, ReMAX1, LeEIX1/2, CuRe1, Ve1/Ve2 and ELR,., activating immunity by recognizing effectors and PAMPs from the phytopathogens (Jones *et al*., 1994; Kawchuk *et al*., 2001; Ron & Avni, 2004; Jehle *et al*., 2013; Du *et al*., 2015; Hegenauer *et al*., 2020; De La Rosa *et al*., 2023; Dixon *et al*.).

In order to understand the distribution and diversity of different receptors across the genome, population, and at the species scale, first, we need a reliable and reproducible bioinformatic tool to annotate complete gene families with high accuracy. Recent studies have mainly devised methods to annotate NLRs, paving the way to systematically understand their distribution and diversity (Steuernagel *et al*., 2020; Ngou *et al*., 2022b, 2024). A few pipelines were devised to annotate PRRs, but either they have a few drawbacks or are not entirely reproducible.

Trying to establish a clear-cut division among RLPs, our recent work (Steidele & Stam, 2021) classified the known RLPs in *A.thaliana* into defense and development-related clusters, phylogenetically. Further highlighted them by functionality, their transcriptomic, and proteomic signatures under different biotic and abiotic conditions. However, here, out of 57 annotated RLPs in *A.thaliana*, 20 were considered as development-related, yet a couple of well-known defence-related RLPs like ATRLP1 and ATRLP3 were found inside the developmental cluster, leaving a gap to improvise the phylogenetic clustering. Another study has identified C2 (Island domain) as conserved region in RLPs, that might shape defence and developmental RLPs, which might help to resolve the precise splitting of RLPs, phylogenetically (Snoeck et al., 2023).

Whereas several studies have focused on predicting RLPs and RLKs in plants to study their diversity, expression, and evolutionary patterns. However, the associated annotation pipelines are either not reproducible or not freely accessible, or they fail to distinguish RLKs with discontinuous kinases from RLPs. Here, we presented an improved pipeline, PlantLRR-PRR, for predicting genome-wide distributed putative RLPs and RLKs. Utilizing the outputs, we examined the inter- and intra-specific RLP diversity in wild tomato species. Most importantly, we tried to dissect the process of their dynamic variation by exploring the phylogenetic grouping of putative defense- and development-related RLP clusters, their conservation, local expansion, and pseudogenization between *Solanum spp* and below we discussed them in detail.

## Methods

### 5’ upstream extension and validation of gene sequence with a new translation initiation start site

Preliminary RLP/RLK annotations revealed a truncated N-terminal or missing SP in a couple of conserved proteins, like Too-Many-Mouths (TMM) (RLP) in the *Solanum pennellii* and *Solanum lycopersicoides* annotations (Bolger et al., 2014; Muir et al., 2014; Powell et al., 2022). To resolve this issue, we developed a simple pipeline that extends 5’ upstream of all gene sequences and validated with the correct start codon (translation initiation start site=TIS). First, 5’ upstream of all genes, including their coding and mRNA sequences, were extended, and extended proteins were extracted and used as a query. We BLAST them against the non-redundant protein database downloaded from NCBI (Camacho et al., 2009) for the respective plant families, such as *Solanaceae* for tomato proteins, Brassicaceae and Fabaceae for *Arabidopsis* and common bean proteins (Camacho et al., 2009; The UniProt Consortium et al., 2025). Blast hits with >85% coverage and identity were taken as positive for individual queries and further considered and checked for the transcriptomic signatures in their extended mRNA regions. Here, the transcriptomic data were collected from multiple sources for the exact individual genotypes using the sra-tools kit [3.2.1] (Supplementary data 1). Next, RNA reads were quality checked with multiQC v1.28 (Ewels et al., 2016). Later, with the TrimGalore tool v0.6.10, the bases with a PHREAD score of <5 and adaptor sequences were trimmed, and reads with <20 bp were removed to select better-quality reads (Krueger, 2023). At last, high-quality RNA reads were aligned to the extended mRNA sequences using HISAT2 v2.2.1 (Kim, D., Paggi, J.M., Park, C. et al., 2019), and merged all BAM files using Samtools v1.19.1 (Li *et al*., 2009). The aligned reads were manually visualized in IGV v2.16.2 (Robinson, 2011) to find the transcriptomic signatures in the extended region. Finally, using ATGpr (web-based), a Kozak motif identification tool, we identified potential Kozak motifs TIS in the extended mRNA/cdna sequences (Kozak, 1978) (Supplementary data 6). After obtaining confirmatory results from BLAST analysis, transcriptomic signatures, and a potential TIS site for the extended sequences, we updated the original GFF file using these findings and utilized the updated proteome for RLP/RLK annotations. Otherwise, we maintain the originals.

### Construction and fine-tuning of HMMs of the PlantLRR-PRR pipeline

To construct the core HMMs for the pipeline, we first extracted putative known RLPs, i.e., 53 from Arabidopsis, 45 from Tomato, and 17 functionally characterized RLPs from Tomato and *Arabidopsis* (Wang *et al*., 2008; Wu *et al*., 2016; Kang & Yeom, 2018; Steidele & Stam, 2021; The UniProt Consortium *et al*., 2025) and aligned them using MUSCLE5 (Edgar, 2004). From aligned sequences, the C2-C3F region (C2=Island domain and C3F=last 4 LRRs till Tryptophan and phenylalanine amino acid in the F domain or TMD) (Steidele & Stam, 2021; Snoeck *et al*., 2023; Ngou *et al*., 2024) was extracted (Supplementary data 5) and a C2C3F-HMM profile was built using the HMMER 3.1 b2 (Eddy, 2023). Next, an LRR-HMM (for extracting LRR-containing proteins), a Pkinase-HMM (for extracting kinase proteins), and an NLR-HMM (for segregating and removing all NLR proteins) profile were constructed using their respective protein domain information obtained from the Pfam database (Supplementary Table 2). Finally, four HMMs were assessed to find a suitable e-value cutoff. For this, we fed the most recent *A. thaliana* primary proteome (Cheng et al., 2017) to individual HMMs and collected protein hits for different e-values and validated by matching against known 57/223 RLPs/RLKs gene IDs from *Arabidopsis* (Wang et al., 2008; Wu et al., 2016).

### Reading frame of the PlantLRR-PRR pipeline

In the first-half of PlantLRR-PRR, HMMs are interconnected. Here, the output of the previous HMM serve as the raw input to the next. Initially, the primary transcript proteome is fed to the C2C3F-HMM, and hits with signatures of the last 4 conserved LRRs (Ngou et al., 2024) are collected. This output is fed to LRR-HMM, collecting all proteins with LRR-domains, and transferred to Pkinase-HMM to segregate all crude RLKs. The remaining proteins are passed to NLR-HMM to find and remove all NLR proteins. However, a few proteins with discontinuous kinases were undetected during the initial crude RLKs filtering. Therefore, all the remaining non-NLR proteins are scanned with Interproscan5 using flags for SuperFamily and CATH-Gene3d databases (Jones *et al*., 2014). All hits with a discontinuous kinase domain are added to the collection of crude RLKs. At last, remaining hits from Interproscan analysis are classified as crude RLPs. In the second half, crude RLPs and RLKs are further classified as full-length RLPs and RLKs based on the presence/absence of SP and TMD. The web-based DeepTMHMM v1.0.44 (Hallgren et al., 2022) and locally installed SignalP4.1 (Nielsen & Kihara, 2017) annotates SPs. DeepTMHMM v1.0.44 (Web-based), Phobious v1.01 (locally installed), and NetGPI 1.1 (web-based) software annotated TMDs (Käll *et al*., 2004; Gíslason *et al*., 2021; Hallgren *et al*., 2022). From here onwards, we designate the full length (having both SP and TMD) of RLPs and RLKs as simply RLPs and RLKs, The complete pipeline can be found on Github (https://github.com/PHYTOPatCAU/RLP_identification)

### Benchmarking the PlantLRR-PRR pipeline using proteomes from BUSCO-validated genomes

We assessed all the genomes used in this study for quality and completeness using BUSCO5.7.1 (Manni et al., 2021) (Supplementary Table 1) with the following settings: “busco -i <genome.fa> -l <lineage.db> --offline --download_path busco_downloads -o <output_file_path> -m genome -f”. To benchmark our PlantLRR-PRRs pipeline, we extracted genome-wide identified RLPs/RLKs from 5 different plant species and compared against the results from previous annotation study (Ngou *et al*., 2022b). We used proteomes with primary gene models from *A. thaliana, A. halleri, P. vulgaris, S. lycopersicum,* and *S. tuberosum* for benchmarking (Supplementary Table 1).

### Multiple sequence alignment and phylogenetic analyses

It is less-likely to get a high-quality MSA for the complete lengths of genome-wide annotated RLPs, as LRR domains are modular in nature and often yield mismatches during sequence alignments. As a result, only the regions around the TMD and its preceding LRRs across the C2 region were extracted and subjected to MSA with MAFFT v7.490 (Katoh & Standley, 2013). We added 16 reference RLPs with putative functionalities, i.e., eight known developmental from *A. thaliana* (RLP4, RLP10/CLV2, RLP17/TMM, RLP29, RLP44, RLP51, RLP55, RLP57) (Fritz-Laylin et al., 2005; Steidele & Stam, 2021) and eight known defense-related RLPs reported in *Solanum spp* for various disease resistance (Cf9, Cf5, EiX1, EiX2, Ve1, Ve2, ELR and CuRe1) (Jones *et al*., 1994; Kawchuk *et al*., 2001; Ron & Avni, 2004; Du *et al*., 2015; Hegenauer *et al*., 2020; Dixon *et al*.). We extracted the C2-C3F region from MSA and constructed a phylogenetic tree with RaxML-HYBRID-AVX (Stamatakis, 2014) with the PROTCATGTR model with 1000 bootstraps (mpirun raxml-HYBRID-AVX -f a -x -p -N -m PROTCATGTR -# 1000 -s ${input} -n {output}), similar to previous study (Steidele & Stam, 2021).

We constructed the complete LRR-PRR phylogenetic tree using crude RLPs and RLKs concatenated from all 8 *Solanum* genotypes. First, we performed MSA for whole proteins with MAFFT v7.490 (mafft –auto input.fast > mafft-output.fasta (Katoh & Standley, 2013) for faster alignment of ∼3600 protein sequences). Then, a complete phylogenetic Tree was constructed with FastTree (Price et al., 2009) (FastTree -wag input.fasta > output.Tree).

We constructed a species tree based on orthologs of the Cure1 cluster shared across five *Solanum spp.* Each set of shared orthologs was separately aligned at the full-length protein level using the default MAFFT (Katoh & Standley, 2013). Regions with gaps were trimmed, and partition files were produced with “Trimal” and the AMAS (Alignment manipulation and summary) tool, respectively (Capella-Gutiérrez *et al*., 2009; Borowiec, 2016). Later, individual gene trees were constructed with IQ-Tree2 (Minh et al., 2020) using 1000 bootstrap replicates, and the resulting four individual gene trees were concatenated. Finally, with ASTRAL (Mirarab et al., 2014), a species tree was generated.

### Orthogrouping of known RLPs in Solanum spp

We identified the size of orthogroups (orthologs and recent paralogs) for the each 8 known defense *(Solanum spp.)* and developmental (*Arabidopsis thaliana*)-related RLPs (Jones *et al*., 1994; Kawchuk *et al*., 2001; Ron & Avni, 2004; Du *et al*., 2015; Hegenauer *et al*., 2020; Steidele & Stam, 2021; Dixon *et al*.) using Blast analysis (Sonnhammer & Koonin, 2002; Camacho *et al*., 2009; Kang & Yeom, 2018). Here Blastp was performed with 16 reference RLPs against the whole primary proteome of all 8 individual *Solanum* genotypes (Supplementary table 4), with an e-value threshold of 1e-10 and a coverage >= 70% (Enright, 2002; Rameneni *et al*., 2015; Yang *et al*., 2020; Pei *et al*., 2021; Li *et al*., 2021). From the output, we segregated the number of BLAST hits for the respective query sequences starting from 30% to 95% sequence identity with increments of 5% and depicted them as a graph. From the graph, we identified an elbow point (a percentage identity threshold displaying a sudden dip in the number of hits relative to all thresholds, red line), beyond the elbow point, we calculated percentage (of number of hits) loss for the difference between two consecutive percentage identity thresholds and identified the threshold recording the lowest loss (Supplementary data 7). Based on this, we set an identity threshold of >=55% and >=45% for identifying size of orthogroups among the known defense- and developmental-related RLPs, respectively (Supplementary data 7).

### Testing Crip21 response among Solanum spp

We tested the response of Crip21 (a *Cuscuta reflexa* elicitor) (Hegenauer et al., 2020) among *S. lycopersicum, S. pennellii,* and *S. chilense* (these 3 genotypes were maintained under optimal conditions in our greenhouse .i.e, 24 °C, 16 h of light). Crip21 peptides were synthesized by Genscript USA Inc. A stock of 10 mM was prepared by dissolving in DMSO, and a working solution of 1 μM was prepared by diluting in sterile water. 1 μM of peptides and water as a negative control was infiltrated into the abaxial side of target leaves at least with three repetitions, and the corresponding response was recorded after 5 days.

### Annotation of Transposable Elements and estimating the proximity of the first nearby TE to each annotated LRR-RLPs

We annotated and extracted the genomic features of de novo-annotated genome-wide distributed transposable elements using the “Extensive de novo transposable elements annotation tool” (EDTA) (Ou et al., 2019). EDTA was run on all the individual unmasked genomes from all 8 *Solanum* genotypes used in this study. The proximity of the first nearby TE in relation to annotated LRR-RLPs was extracted using bedtools (Quinlan & Hall, 2010). First, we sorted the GFF file, “bedtools sort -i inputgff3 > input_sorted.gff3”, and later extracted the distance of the closest TEs using “bedtools closest -a EDTA.gff -b input_sorted.gff3 -d > TE_with_distance.txt” and finally extracted the distance for our target gene family.

## Results

### Optimisation of the PlantLRR-PRR pipeline

Our RLP/RLK annotation pipeline, PlantLRR-PRR, builds on the earlier examples that both RLP and RLKs share the conserved last 4 LRRs (also known as the C3 domain) as defining features (Ngou et al., 2024). First, to standardize HMMs’ sensitivity to capture crude RLPs and RLKs, primary transcript proteomes were scanned individually with C2C3F-HMM, LRR-HMM, and Pkinase-HMM under inclusive thresholds, ensuring that no true candidates were missed. Then, subsequent filtering (using an NLR HMM) removed NLR proteins from the dataset and separated crude RLPs from RLKs with high confidence (Figure 1A).

**Figure 1:**
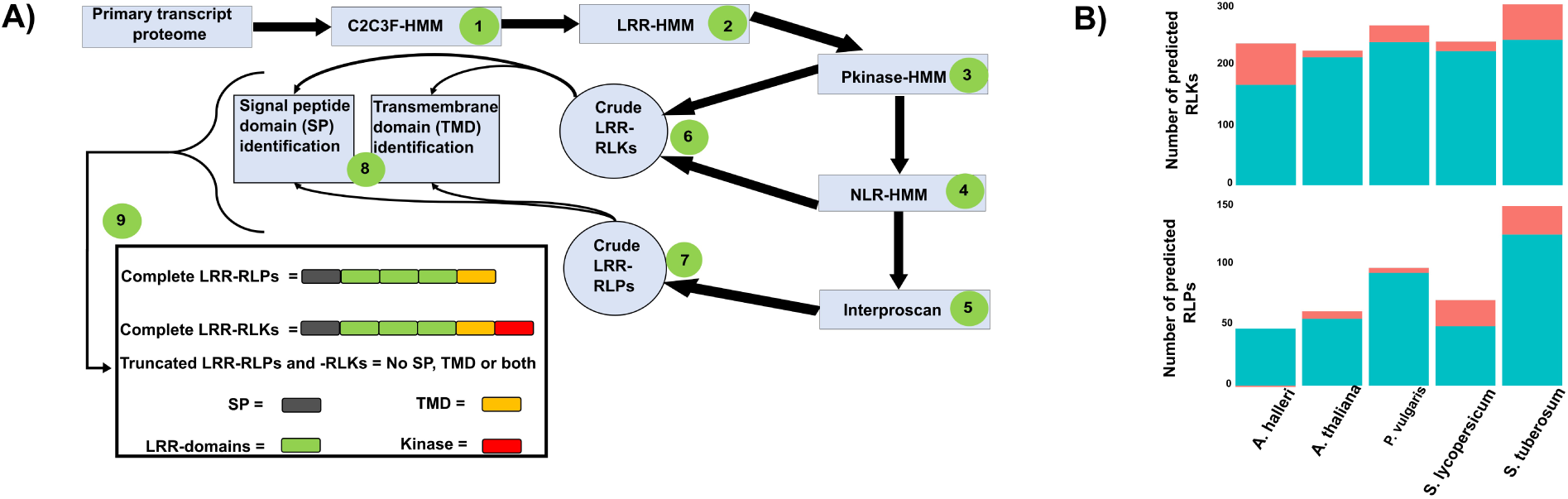
PlantLRR-PRR pipeline and comparison of predicted RLPs and RLKs among five different plant species against results from the previous study. A) Outline of the Plant LRR-PRR pipeline. B) Visualisation of predicted, genome-wide distributed complete versions of RLPs/RLKs from five different plant species in the current analysis compared to the previous analysis by Ngou et al., 2022. The Teal part of the stacked bar charts represents the RLPs/RLKs count annotated in Ngou et al., 2022, which also overlapped with our results. The light red part of the stacked bar charts represents the additional positive and negative counts of RLPs/RLKs annotated in our analysis.

We fine-tuned the e-value threshold of each HMM by testing it with the *Arabidopsis thaliana* proteome, Araport11 (Cheng et al., 2017) under variable e-values (1E-1 to 1E-20). Depending on the threshold, the models retrieved between 325–487 C2-C3F hits, 163–767 LRR hits, 999–1055 kinase hits, and 216–801 NLR hits from C2C3F-HMM, LRR-HMM, Pkinase-HMM, and NLR-HMM, respectively (Supplementary data 2). To identify an optimal cutoff, we compared our outputs with 57 and 223 annotated putative RLPs and RLKs from *Arabidopsis* (Fritz-Laylin *et al*., 2005; Wu *et al*., 2016; Steidele & Stam, 2021). With 1E-3 and higher, we retrieved 54 of 57 previously annotated RLPs and 222 of 223 RLKs annotated from *A. thaliana* (Fritz-Laylin *et al*., 2005; Wu *et al*., 2016; Cheng *et al*., 2017; Steidele & Stam, 2021) (Supplementary Figure 1 and Supplementary data 2). The three missing RLPs (AT4G13900, AT1G54480, and AT2G15040) were absent in this version of the proteome of *Arabidopsis* (Cheng et al., 2017). Interestingly, the gene AT3G46350, previously annotated as RLK (Wu et al., 2016), was rightly annotated as a crude potential Malectine-like RLP. Thus, by maintaining an evalue of 1E-3 for all HMMs, the pipeline has picked up all known RLPs and RLKs.

### PlantLRR-PRRs pipeline identifies additional RLPs and RLKs in *A. thaliana* and in other model species

Running the interconnected HMMs in the PlantLRR-PRR pipeline on the published *A. thaliana* primary transcript proteome (Cheng et al., 2017) yielded a comprehensive set of 135 and 228 crude RLPs and RLKs for downstream annotation (Supplementary data 2). Annotating the crude versions for SP and TMD resulted in 59 and 223 full-length RLPs and RLKs (Figure 1B). Of the 59, 50 overlap with 57 previously annotated RLPs (Fritz-Laylin et al., 2005; Wang et al., 2008); three proteins (AT1G54480, AT2G15040, AT4G13900) were absent from the updated proteome (Cheng et al., 2017). The remaining four, AT1G74200 and AT2G33030 lacked SP; AT2G33080 lacked a TMD; while AT3G53240 lacked both TMD and SP. Additional we annotated nine RLPs, including two Malectin-domain-having RLPs: AT3G46350, previously mis-annotated as RLK (Wu et al., 2016) and AT1G25570; and 7 novel RLPs: AT1G54470, AT2G15042, AT3G25670, AT4G09435, AT4G28560, AT4G16162, and AT5G49750. Interestingly, we found one additional RLP, AT1G33590.2 (Supplementary figure 2), and an RLK, AT4G20940.1, due to a successful 5’ upstream extension of the *A. thaliana* genes. The final dataset included 60 and 224 RLPs and RLKs, respectively (Figure 1B and Supplementary data 2).

We tested the pipeline on an additional four primary proteomes from *A. halleri*, *P. vulgaris, S. lycopersicum,* and *S. tuberosum* (Supplementary Table 1). First, their genome completeness was assessed using BUSCO 5.7.1 (Manni et al., 2021). They were in the range of 99.1–99.6% (Supplementary Table 2). Next, we compared our annotation outcomes with previous studies (Ngou *et al*., 2022b), and our pipeline predicted a higher number of RLPs and RLKs in all four species (Figure 1B) (Supplementary Tables 3–4). RLKs were increased by 69 in *A. halleri*, 28 in *P. vulgaris*, 16 in *S. lycopersicum*, and 59 in *S. tuberosum*. Further, RLPs were increased by 4 in *P. vulgaris,* 21 in S*. lycopersicum*, and 23 in *S. tuberosum* (Supplementary Table 3). Manual inspection of some cases highlights the robustness of the pipeline: “Ah1G34300.3” in *A. halleri* was mispredicted as an RLP (Figure 1B), but with the inclusion of the interproscan analysis, it is annotated as an RLK with a discontinuous kinase domain. Thus, our improved pipeline enables standardized (re)classification of RLPs and RLKs.

### Copy number variation of RLPs and RLKs, and chromosomal distribution of RLPs in *Solanum spp*. of the tomato clade

To investigate Copy Number Variation (CNV) in the RLP and RLK families within and between tomato species, we applied the PlantLRR-PRR pipeline on eight tomato genotypes belonging to 5 different *Solanum* species (Supplementary Table 4). BUSCO completeness for each genome ranged from 99.1% to 99.4% (Supplementary Table 1).

First, RLP/RLK interspecies CNV was assessed from five genotypes representing five tomato species;*S. lycopersicum, S. pimpinellifolium, S. chilense, S. pennellii,* and *S. lycopersicoides.* We found moderate and low levels of relative RLPs and RLKs CNV, with 22% and 4% of the coefficient of variation, respectively (Supplementary Figure 3B). RLP count ranged between 60 (*S.pennellii*) and 94 (*S.lycopersicoides).* Whereas RLK count varied from 232 (*S.pennellii)* to 258 (*S.lycopersicoides)* (Figure 2A). We also confirmed that these counts show only minor deviation in the Q-Q plots, and a Shapiro-Wilk test returned a “W” value close to 1 with p > 0.05; therefore, we conclude that there are no obvious outliers or sequencing and genome assembly artefacts (Supplementary Figures 3A-B) and that this level of CNV represents true biological diversity.

**Figure 2:**
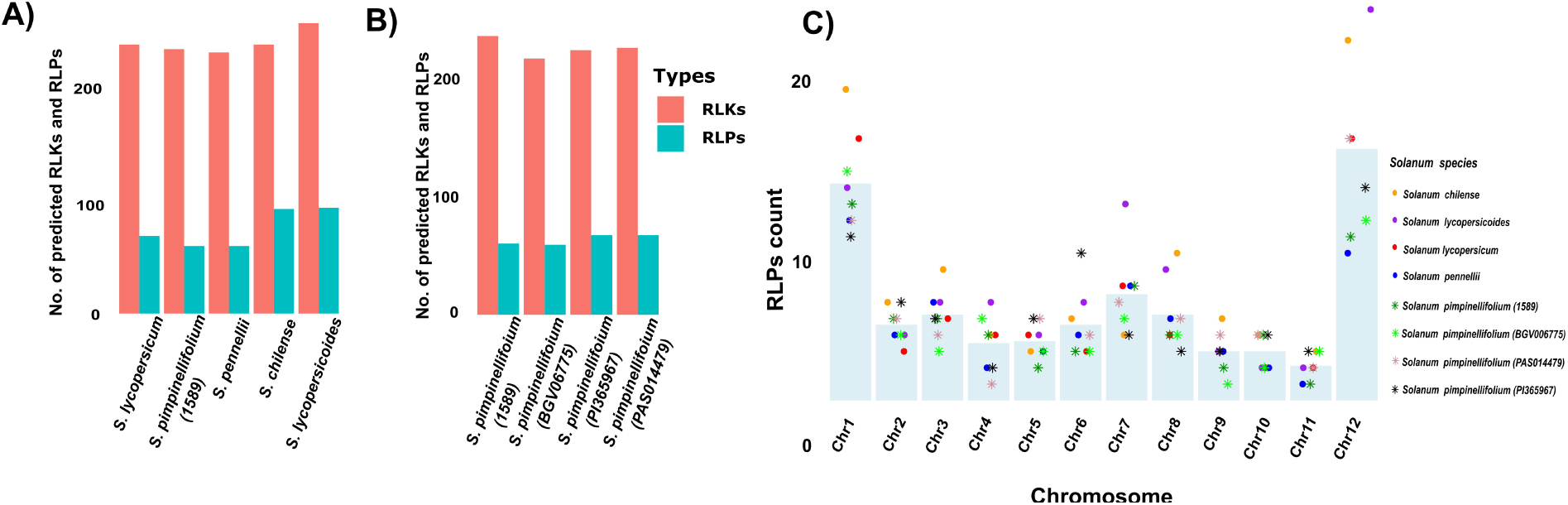
Representation of the CNV of RLPs and RLKs, and chromosomal distribution of RLPs between and within different genotypes of *Solanum spp*. A-B) Visualisation of CNV of genome-wide extracted RLPS/RLKs between five *Solanum species* (A) and between four genotypes of *S. pimpinellifolium* (B). The X and Y axes represent individual species and RLPs/RLKs gene counts, respectively. Orange and teal bars represent RLKs and RLPs counts for each studied genome, respectively. C) Visualisation of the chromosomal distribution of RLPs counts across each of the eight *Solanum* genotypes. The X- and Y-axis represent the chromosome numbers and RLPs counts, respectively. Each coloured dot on the respective chromosome depicts the RLPs extracted from that chromosome from one of the eight *Solanum* genotypes. The tip of individual barplots depicts the mean count of RLPs on the respective chromosomes.

Next, we dissect the RLP and RLK CNV within one *Solanum spp* with four different genotypes of *S.pimpinellifolium* (1589, BGV006775, PAS014479, and PI365967) (Supplementary Table 4). We found relatively lower CNVs for both RLPs and RLKs with 7% and 3% coefficient of variation, respectively (Supplementary Figure 3D). Here, RLP counts ranged from 59 to 67, and for RLK from 216 to 235 (Figure 2B, Supplementary Table 4). Again, Q-Q and a Shapiro-Wilk test showed no evidence for outliers or sequencing artefacts (Supplementary Figure 3C-D). Thus, we conclude that indeed there is CNV for RLPs and RLKs within the species.

Overall, RLK copy numbers might appear more stable both between and within *Solanum spp.* However, RLPs were more fluctuating. Additionally, RLPs are better known to have specific functions in either defence or developmental activities, so we will focus on the RLP gene family in the remainder of the manuscript.

Focusing on the RLP family, we asked whether RLP distribution is constrained to a few chromosomal locations. So, we visualised the distribution of RLPs across the 12 chromosomes (12 Chr) (Figure 2C) in all eight *Solanum* genotypes. Primarily, RLPs are found on each chromosome irrespective of the genomes. Across all eight genotypes, most of the RLPs (∼40%) accumulated on Chr1 and Chr12, and the least (∼3%) on Chr11. A chi-squared test revealed that *S. pennellii (0716)* and *S. pimpinellifolium (1589)* have an even distribution of RLPs across all chromosomes (P>0.05), whereas the remaining six *Solanum* genotypes showed uneven distributions. Further, we observed species-specific chromosomal RLP hotspots. E.g., *S. lycopersicoides* (light-purple dot) and *S. chilense* (light orange) accessions both have almost double the average number of RLPs on chromosomes 12 and 8; *S. lycopersicoides* alone has twice the average number of RLPs on chromosome 7, but *S. chilense* has less than half of that. *S. pennellii* always has about the average number of RLP or less, identified on each chromosome.

### Defence associated RLPs are highly variable between species

Next, we sought to categorize RLPs into putative defense and developmental types phylogenetically. First, we improved the phylogenetic classification of *A.thaliana’s* RLPs into defense and developmental superclades (see methods). This resulted in a clear improvement compared to the previous study (Steidele & Stam, 2021) with two superclades. All biotic-stress-induced, transcriptionally active RLPs (including AtRLP1 and AtRLP3) (Steidele & Stam, 2021) grouped into the defense clade. The 8 and 6 known (highlighted in blue and red colours) known putative developmental and defense-related RLPs (Steidele & Stam, 2021) were clearly segregated and distributed inside respective superclades (Supplementary Figure 4).

Using this methods, we classified RLPs in five *Solanum spp*, by construting individual phylogenetic trees of their predicted RLPs and included 16 reference RLPs. Resulting phylogenetic trees consistently resolved into two superclades, segregating defence (red arc) and developmental clusters (green arc) (Supplementary Figure 5) and all reference genes fit where expected. This revealed that *S. pennellii* and *S. pimpinellifolium* (1589) contained a relative proportion of 72% and 75%, whereas *S. lycopersicum*, *S. lycopersicoides*, and *S. chilense* had 80–82% of putative defence-related RLPs compared to all extracted RLPs from respective genotypes. To provide a comparative framework, we constructed a composite phylogenetic tree that included all predicted RLPs from the five *Solanum species* and 16 reference RLPs (Figure 3A). The phylogeny robustly split into two superclades, i.e., defense clade: red arc, and developmental clade: green arc, with 100% bootstrap support; differences in size of the defence associated RLPs can be observed. Next, we constructed species-wide known RLP orthogroups, represented as heatmaps (Figure 3B-C). These reveal that the elevated fraction of defense RLPs in *S. chilense* and *S. lycopersicoides* results from increased numbers in the Cf9 and ELR clusters with 9-12 and 12-14 copies, respectively (Figure 3A-B). *S. pimpinellifolium* had only 5 and 7 copies for Cf9 and ELR clusters, respectively (Figure 3B). We also found that the CuRe1 cluster is compromised in *S. pennelii*, with only one potential ortholog, compared to a minimum of 4 in the remaining species (Figure 3B). We also found that Ve2 is highly conserved among all the *Solanum spp* with >95% identity; this in contrast to Ve1 which fits the observation that Ve1 is the decoys of Ve2 (Kawchuk *et al*., 2001). On the contarary we checked whether Ve2 is conserved in other related *Solanum* species like *Capsicum annum* (Kim *et al*., 2014). To our surprise Ve2 is ∼81% conserved both in-terms of coverage and percentage identity.

**Figure 3:**
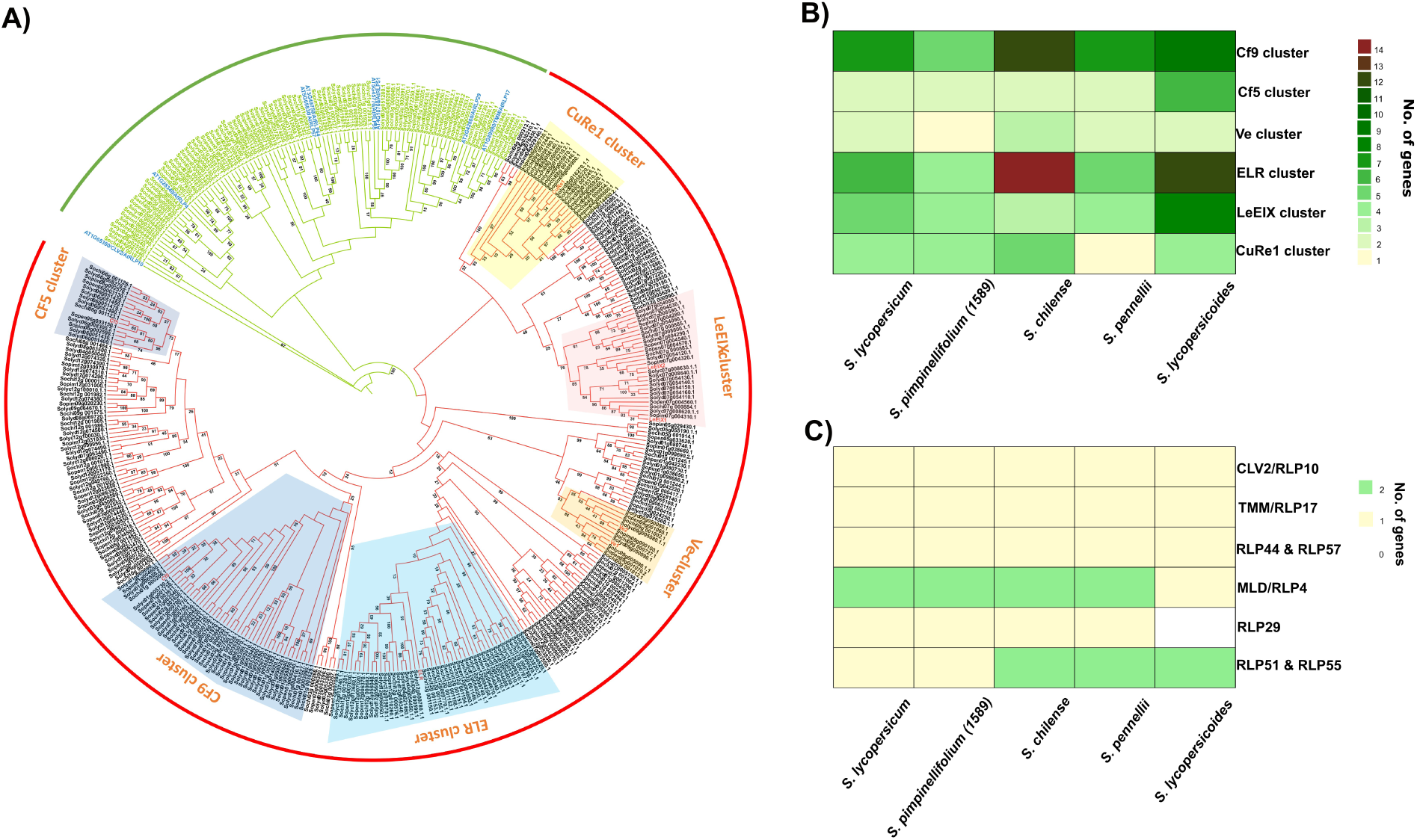
Phylogenetic inferences and orthogroups of extracted RLPs from 5 *Solanum spp.* with 16 reference RLPs. A) A combined phylogenetic tree was constructed targeting the C2C3F domain from all the predicted RLPs of five *Solanum spp*. Sixteen RLPs with established functionalities were included as a reference. Eight Arabidopsis developmental RLPs: RLP17/TMM, 51, 55, 29, 4, 10/CLV2, 44, and 57 (blue labeled) and eight Solanum defense-related RLPs: Cf9, Cf5, ELR, Ve1, Ve2, EiX1, EiX2, and CuRe1 (red labeled). The tree was inferred with RAxML using 1,000 bootstrap replicates. Two well-supported superclades, similar to Supplementary figure-1 (100% bootstrap strength), corresponding to developmental and defense-related RLPs were indicated by green and red arcs, respectively. B-C) Heatmap showing the size of the orthogroups identified for the known defense-(B) and development-related genes(C). Gradient colors represent RLPs counts ranging from white(lowest) to brown (highest).

As expected, we found development-associated RLPs to be more conserved. TMM and CLV2 were maintained as single-copy orthologs in all five *Solanum spp* (Figure 3C). But, there were a few anomalies: RLP44 and 57, which are paralogs in *A. thaliana* (Wang *et al*., 2008; Steidele & Stam, 2021), are represented by a single gene in all the *Solanum spp.* RLP29 was missing in *S. lycopersicoides* (devoid of SP). RLP4 has two paralogs in all *Solanum spp* except in *S. lycopersicoides.* And the RLP51 and RLP55 orthogroup have only one gene in *S. lycopersicum and S. pimpinellifolium (1589)* (Figure 3C).

### Elevated intraspecific stability of RLP clades might support functional conservation in *S. pimpinellifolium*

Next, we reconstructed individual phylogenetic trees for the four *S. pimpinellifolium* accessions and constructed RLP orthogroups by including all 16 reference RLPs. (Figure 4A-D). As anticipated, the phylogenetic tree was split into defence and developmental superclades (Figure 4A). In all four genotypes, ∼25-30% and 70-75% relative proportion of RLPs clustered with developmental and defense-related phylogenetic superclades, respectively. The Cf9, Cf5, and ELR clusters show strong variation, ranging from 4-8, 1-7, and 2-7, respectively, between accessions. The Ve cluster had only one copy in both 1589 and BGV006675 genotypes, whereas two copies were found in all other genotypes (Figure 4E) and across *Solanum spp* (Figure 4B). Also, here Ve2 is more conserved than to Ve1 in all 4 accessions. The CuRe1 cluster had 2 copies in PAS014479 and PI365967 compared with four copies in the other genotypes (Figure 4E) and most other *Solanum spp.* (Figure 4B). Development-related orthogroups were much more stable within the species. Diversity was only found for the RLP51 & RLP55; there was one copy lacking from the 1589 genotype (Figure 4F).

**Figure 4:**
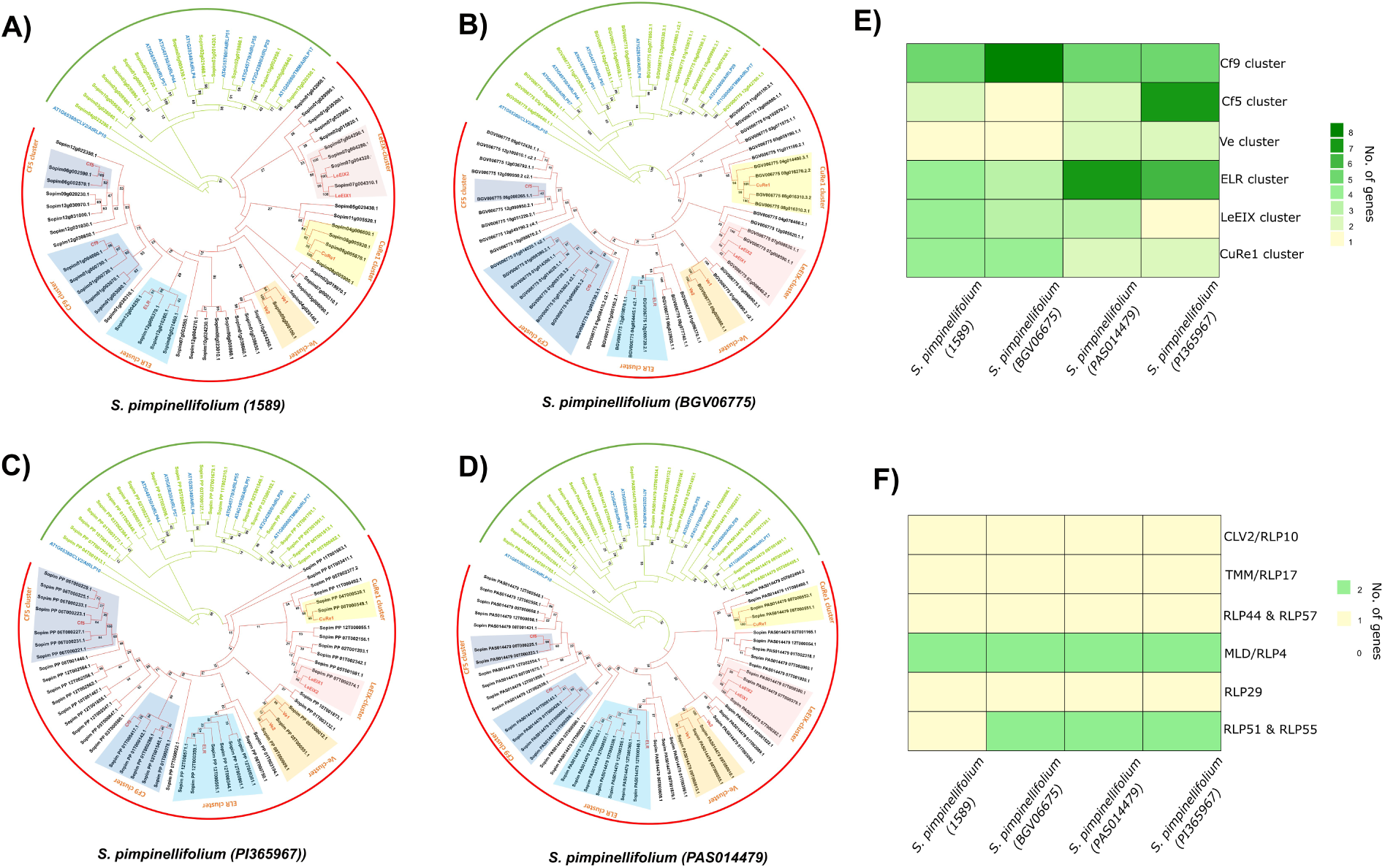
Phylogenetic inferences and orthogroups of RLPs from 4 *S. pimpinellifolium* genotypes clustered with 16 reference RLPs. A-D) Individual phylogenetic trees were constructed targeting the C2C3F domain from all the predicted RLPs of four *S. pimpinellifolium* genotypes (A. SP-1589, B. SP-BGV006775, C. SP PI365967, D. SP-PAS014479). Sixteen RLPs with established functionalities were included as a reference. Eight Arabidopsis developmental RLPs: RLP17/TMM, 51, 55, 29, 4, 10/CLV2, 44, and 57 (blue labeled) and eight Solanum defense-related RLPs: Cf9, Cf5, ELR, Ve1, Ve2, EiX1, EiX2, and CuRe1 (red labeled). The tree was inferred with RAxML using 1,000 bootstrap replicates. Two well-supported superclades, similar to Supplementary figure-1 (100% bootstrap strength), corresponding to developmental and defense-related RLPs were indicated by green and red arcs, respectively. E-F) Heatmap showing the number of orthologs and recent paralogs identified for the known defense-related genes (E) and development related genes(F). Gradient colors represent RLPs counts ranging from white(lowest) to brown (highest).

### Lineage-specific expansion and pseudogenization of RLP cluster

To validate the observed cluster size differences at the species level, we examined the genomic organisation of the CuRe1 and Cf5 loci among *Solanum spp.* In all 5 *Solanum spp.,* the CuRe1 cluster contains at least 4 paralogs (Supplementary Figure 5). However, in *S. pennelli,* only a single putative functional CuRe1 paralog was annotated, and the remaining three were pseudogenized and annotated as truncated RLPs. To confirm the pseudogenization, orthologs of all four CuRe1 members were extracted from each species, aligned, and used to construct a species tree (Figure 5A, Supplementary Figure 6). As expected, the generated species tree (Figure 5A) depicts the evolutionary relationship of *Solanum spp* (Pease *et al*., 2016). Inspection of the alignments confirmed pseudogenization, revealing loss of the SP in Sopen04g006530 and loss of the TMD in Sopen08g006560 and Sopen08g006740 (Supplementary Figure 6). The remaining CuRe1 paralog, Sopen08g006730, contained with a retrotransposon insertion, suggesting that it may also be non-functional. To test whether these structural changes affected receptor function, we infiltrated Crip21 into *S. lycopersicum*, *S. chilense*, and *S. pennellii*. Whereas *S. lycopersicum* and *S. chilense* developed a clear cell-death response, *S. pennellii* showed no response (Figure 5A), consistent with loss of a functional CuRe1 receptor (Figure 5A) (Hegenauer *et al*., 2020).

**Figure 5:**
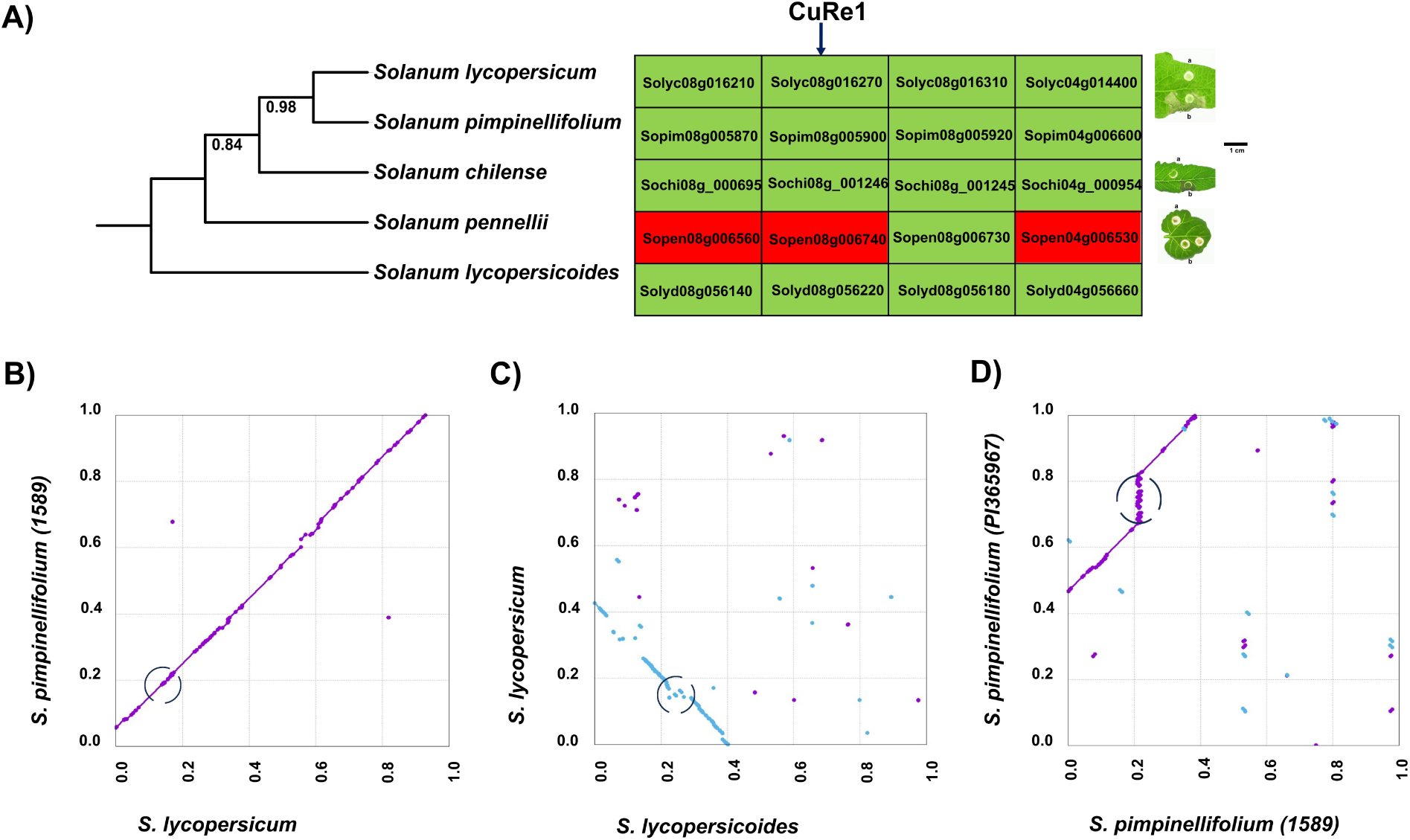
Pseudogenization of CuRe1 cluster in *S. pennelii* and radiation of Cf5 cluster between and within *Solanum spp.* A) Visualisation of species tree among 5 *Solanum* species utilizing orthogroups of CuRe1 cluster and displaying the pseudogenization of the CuRe1 cluster in *S. pennellii* compared to other *Solanum spp*. Red and green boxes indicates putatively pesudogenized and functional CuRe1 homologs, respectively. B) Dotplot of Cf5 loci showing balanced synteny at genomic level between *Solanum spp* i.e., *S. pimpineliifolium* 1589 and *S. lycopersicum.* C) Dotplot of Cf5 loci showing syntenic diferrnces at genomic levels between *S.lycopersicoides* and *S. lycopersicum*, at species level. D) Dotplot of Cf5 loci showing syntenic diferrnces at genomic levels between *S. pimpinellifolium* (PI365967) and *S. pimpineliifolium* 1589, within species level.

In contrast to the contraction of the CuRe1 locus, the Cf5 cluster showed lineage-specific expansion among *Solanum* species. To confirm that the observed expansion is true and not an annotation error, we generated dotplots focusing on Cf5 loci at the genomic level. Dot plots showed conserved locus size between *S. lycopersicum* and *S. pimpinellifolium* (1589) (Figure 5B), whereas clear expansions were observed in *S. lycopersicoides* (Figure 5C) and *S. pimpinellifolium* PI365967 (Figure 5D).

### Defence-related RLPs are in proximity to transposable elements

In both of the above scenarios, we noticed transposable elements as a probable driving force for these dynamics. In the first case, the only putative-active gene in the CuRe-1 cluster of *S. pennellii*, Sopen08g006730, was fused with a retro-transposon. In the Cf5 regions of *S. lycopersicoides* and *S. pimpinellifolium* (PI365967) with high copy number, we found multiple transposable elements flanking the respective Cf5 paralogs like Tc1_Mariner_TIR_transposon, helitron, broken repeat_fragment, L1_LINE_retrotransposon, hAT_TIR_transposon, Mutator_TIR_transposon, PIF_Harbinger_TIR_transposon, Copia_LTR_retrotransposon, LTR_retrotransposon, target_site_duplication, and long_terminal_repeat. However, only a few were annotated in non-radiated paralogs in *S. lycopersicum* and *S. pimpinellifolium (1589)* like Tc1_Mariner_TIR_transposon, LTR_retrotransposon, Mutator_TIR_transposon, and helitron.

To assess the potential role of transposable elements (TEs) in RLP evolution, we quantified the proximity of TEs to defence- and developmental-related RLPs. More than 50% of all RLPs directly overlapped with TEs (0 bp distance; Supplementary data 8). To prove this, we fitted a zero-inflated Tweedie model to check the randomness and trueness of fused TEs to defence-related RLPs and performed a chi-square and Fisher’s exact test to check the true dependency and odds ratio of both groups to the proximity of TE. The zero-inflated Tweedie model fits better than the non-zero-inflated model with AIC:3183.45 vs 3203.18, and ΔAIC = 19.73, and confirmed that TEs are more often associated with RLPs than expected randomly (Table 1). The conditional mean distance shows that the non-zero distance TEs are average ∼485bp away from the defence-related members and that distance is not significantly (p-value=0.589) different from developmental-RLPs (Supplementary table 5 and Supplementary Figure 7). Yet, the Zero-inflation component revealed a significant difference in the probability of TEs being inside a defence ordevelopmental RLP, ∼0.734 for defence (odds=exp[1.0176]=2.77) and ∼0.473 for developmental (odds=exp[1.1258]=0.897). This corresponds to odds of TE overlap being approximately 3 fold higher in defence RLPs relative to developmental RLPs. Additionally, the model-free comparison of zeros vs non-zeros distances corroborated the model’s result. Chi-square and Fisher’s exact test revealed there is a dependency of TE proximity to the developmental and defence-gene cluster, and it is not independent (pvalue<0.05). Fisher’s exact test confirmed a significant association (p = 9.19×10⁻⁷) with an odds ratio of 0.327 (95% CI 0.205–0.519), indicating lower odds of TE overlap among development genes. As a confirmation, the odds ratio from Fisher’s exact test complements the zero-inflation model results. A model-independent analysis supported these results. Both the chi-square and Fisher’s exact tests detected a significant association between RLP class and TE overlap (P < 0.05). Fisher’s exact test estimated an odds ratio of 0.327 (95% CI: 0.205–0.519; P = 9.19 × 10⁻⁷), indicating significantly lower odds of TE overlap among developmental than defence-related RLPs.

## Discussion

RLPs and RLKs are major plasma membrane receptors involved in both developmental and defense activities (Jones & Dangl, 2006; Macho & Zipfel, 2014; Wu *et al*., 2016; Steidele & Stam, 2021). Despite the recognized importance of RLPs in tomato resistance breeding, little is known about how these family evolved within and between wild tomato species. Most previous RLP studies have focused on single species or model plants (Wang *et al*., 2008; Kang & Yeom, 2018; Yang *et al*., 2020; Steidele & Stam, 2021), or extensively studied the evolutionary pattern and origin of the RLPs/RLKs across (Ngou *et al*., 2024) the plant kingdom. Detailed studies on inter- and intra-species variation as found for the NLR receptor family (Van De Weyer *et al*., 2019; Sutherland *et al*., 2024; Silva-Arias *et al*., 2025) are lacking for RLPs in tomato. Moreover, previous RLP/RLK annotation pipelines lack reproducibility or are genuinely incomplete (Ngou *et al*., 2022b).

To address these gaps, we developed a compresensive PlantLRR-PRR pipeline for RLP and RLK identification that starts from 5’ upstream extension and validation of the CDS followed by compresensive motif searches in complete predicted proteomes). Utilizing these, we annotated RLPs and RLKsand looked deeper into RLPs in available tomato genomes

### Pre and post PlantLRR-PRR pipeline improvements

An important improvement preceding the development of the PlantLRR-PRR pipeline was identifying the correct start codon using 5’ extension and validation, highlighting the critical role of accurate reference genome annotation for receptor identification. Without this extension, the conserved RLP, TMM appeared N-terminally truncated in *S. pennellii* and *S. lycopersicoides*, despite its known conservation across plant species (Muir *et al*., 2014). Our correction produced the complete gene models in both species and demonstrated that annotation errors can lead to misclassification of receptor proteins. Validation (Supplementary figure 2) of a newly identified *Arabidopsis* RLP, AT1G33590, with an extended 5′ region further supported the robustness of our approach and is corroborated by the literature (Irshad *et al*., 2008; Masi *et al*., 2016). Such annotation challenges are common in genome-wide receptor studies and can significantly affect comparative analyses (Ngou *et al*., 2022b, 2024; Sutherland *et al*., 2024; Silva-Arias *et al*., 2025). The integration of InterProScan domain analysis further improved the pipeline by enabling more accurate separation of RLKs with degenerated kinase domains from true RLPs. Together, these improvements demonstrate that careful curation of reference annotations is essential for reliable genome-wide identification and evolutionary analysis of plant immune receptors.

Using the PlantLRR-PRR pipeline, we identified and classified LRR-RLPs and RLKs and examined copy number variation across tomato species to better understand the evolutionary dynamics of these receptor families. While RLKs were comparably conserved, RLPs displayed more variation between species and (more moderate) variation still remainede within species (Supplementary figure 3), a pattern similar to that reported for NLR immune receptors (Silva-Arias *et al*., 2025; Benoit *et al*., 2025). This supports the idea that diversification of receptor repertoires primarily occurs over evolutionary timescales, possibly driven by host–pathogen co-evolution, whereas within-species stability reflects functional constraints that may arise from the need to maintain signaling robustness while avoiding fitness costs associated with excessive immune activation (Martin & Tate, 2024). Yet, as with NLRs (Stamatakis, 2014; Silva-Arias *et al*., 2025) relatively large CNV can be observed in some subclades of the RLPs. The uneven chromosomal distribution of RLPs further suggests that local duplication, diversification, and pseudogenization have contributed to receptor evolution, consistent with previous findings in tomato and other plant species (Andolfo *et al*., 2013; Yang *et al*., 2020).

### Lineage and genotypic-specific expansion and conservation of RLPs between and within *Solanum spp*

Using a phylogenetic framework based on the well-characterized Arabidopsis RLP repertoire (Steidele & Stam, 2021), we classified *Solanum* RLPs into developmental and defence-related groups, revealing a typical pattern of deep conservation combined with lineage-specific diversification. Not surprisingly, development-associated RLPs such as TMM and CLV2 were consistently maintained as single-copy genes, suggesting strong purifying selection consistent with their central roles in plant development and primary metabolism. Similar evolutionary stability has been described for core receptor components across plant species, supporting the idea that such regulatory receptors are subject to strong functional constraints, such as tight dose dependency, that might not allow gene duplication (Contreras *et al*., 2023; Sutherland *et al*., 2024). Whereas Ve2 was a notable exception, almost all defence-associated RLPs show evolutionary plasticity, with lineage-specific expansions and occasional gene losses. The higher number of defence-related RLPs in some wild species such as *S. chilense* and *S. lycopersicoides,* compared with cultivated relatives or other wild species supports the hypothesis that immune receptor diversification may reflect adaptation to heterogeneous pathogen environments and ot just lrger phylogenetic changes (Torres Ascurra *et al*., 2023; Silva-Arias *et al*., 2025; Kahlon *et al*.). The uneven chromosomal distribution of RLPs in some species further suggests that local tandem duplication and pseudogenization contributed to this diversification in certain species or accessions, consistent with general models of plant NLR evolution (Michelmore & Meyers, 1998; Leister, 2004).

Examples such as the genotype-specific expansion of the Cf-5 cluster and the partial pseudogenization of the CuRe1 cluster in *S. pennellii* illustrate how local gene duplication and gene loss jointly shape receptor repertoires. These contrasting patterns likely reflect the balance between selection for novel recognition specificities and constraints imposed by receptor network stability or the fitness costs associated with immune gene expansion (De La Rosa *et al*., 2023; Kahlon *et al*.). While the expansion of receptor clusters related to Cf-9, ELR and Cf-5 homologs across some *Solanum* species highlights how local gene family expansion may promote the emergence of new pathogen recognition specificities while maintaining core immune functions, the pseudogenization of CuRe1 in *S. pennellii* suggests that RLPs may also be lost when selective pressure from the corresponding parasites is relaxed. At the same time, the conservation of core CuRe1 paralogs across species indicates that essential recognition functions are maintained despite local gene family turnover. Together, these observations are consistent with a birth–death model of plant immune receptor evolution, in which cycles of gene duplication, diversification, and loss shape receptor diversity in response to changing pathogen pressures (Michelmore & Meyers, 1998).

### TE proximity to defence and developmental RLPs highlights the possible role of TEs in the expansion and pseudogenisation of RLPs across Solanum spp

Our analyses further suggest that transposable elements (TEs) may contribute to the dynamic evolution of RLP gene families across *Solanum* species. Defence-related RLPs showed approximately threefold higher odd to be connected to TEs compared with developmental RLPs, supporting a potential role of TEs in promoting local gene expansion, structural variation, and pseudogenization. Similar patterns have been described for NLR immune receptors, which are often located in TE-rich genomic regions associated with high turnover and copy number variation (Sutherland *et al*., 2024; Silva-Arias *et al*., 2025). The higher proportion of TE insertions within defence-related RLPs compared with developmental receptors suggests that that TE activity may preferentially contribute to the diversification of defence-related receptors (Sutherland *et al*., 2024). Case examples such as the repeat-associated expansion of the Cf-5 locus and TE-associated disruption of CuRe1 paralogs further illustrate how TE-rich genomic environments may facilitate both gene birth and gene loss. The lower TE association observed for developmental RLPs such as TMM and CLV2 likely reflects stronger purifying selection, or limited tolerance of TE integration, as structural disruption of these central signalling components would be expected to have severe fitness consequences. In contrast, defence-related receptors may tolerate greater structural variation because they often function as part of more flexible and possibly partly redundant recognition systems; for example, the Cf4, Cf5, and Cf9 loci all contain putative functional paralogs and all three loci defend against a single pathogen. These genes are involved in defence mediated through direct or indirect gene-for-gene interactions, and the pathogen *Cladosporium fulvum (*now *Passalora fulva*) shows high allelic diversity in corresponding Avr genes (Parniske & Jones, 1999; Van Der Hoorn *et al*., 2001; De La Rosa *et al*., 2023; Kahlon *et al*.; Wulff *et al*.). In contrast, the only known defence RLPs which seemingly showed very high conserrvation was Ve2. It was more stable not only across tomato species but also in *Capsicum annum* (Kim *et al*., 2014) as it shares similarity to helper NLRs compared to its flexible partner, Ve1, as it acts as a plausible sensor (Kawchuk *et al*., 2001; Kalischuk *et al*., 2022; Sutherland *et al*., 2024). Here, Ve1 interact directly with the corresponding effectors from *Verticillium dhaliae* (Ave1), and then pass corresponding messages via Ve2 to activate the defense response. This contrast supports the broader concept that plant receptor families evolve under a balance of constraint and innovation, with core signalling components or co-dependent components remaining conserved while peripheral recognition modules or receptors that function on their own diversify (Contreras *et al*., 2023; Sutherland *et al*., 2024). Also, the conservation of Ve2 might have longer storyline as silencing Ve2 ortholog, caRLP264, in *Capsicum annum* (Kim *et al*., 2014; Kang *et al*., 2022) compromised basal immune response against multiple pathosystems like Tobacco mosic virus, *Xanthomonas axonopodis pv. glycines8ra, Ralstonia solanacearum* and *Phytophthora capsici*. This opens new avenues to consider identifying conserved hub RLPs among *Solanum species* to verify their potentiality in maintaining basal immune response against multitude of phytopathogens. Further investigation of Ve2 is needed among domesticated and wild tomatoes to dissect its underestimated plausible positive regulating or inducing role of basal immune response against wide variety of phytopathogens.

Taken together, this work provides robust identification pipeline for RLPs and shows that it can be used to asses RLP diversity in wild and cultivated species or accessions, moving beyond previous single-reference or crop-focused studies, and providing detailed case examples. By integrating improved gene annotation, phylogenetic classification, CNV analysis, and TE landscape context, our results highlight how RLP repertoires are shaped by the interplay between evolutionary constraint and adaptive diversification. The focus of various wild tomato species reveals patterns of receptor expansion and contraction that would remain hidden in domesticated genomes alone and underscores the importance of studying germplasm for understanding immune receptor evolution (Penczykowski et al., 2026). Hence our pipeline and initial findings open new avenues for linking receptor evolution with ecological adaptation, and functional immune diversity, and provide a foundation for future studies connecting receptor variation to phenotypic resistance and translational applications in crop improvement.

## Author Contributions

NJ designed and conducted the experiments. NJ developed and integrated the software tools and analytical approaches used in this study. NJ performed the data analyses and visualisation. NJ and RS conceived the study and wrote the manuscript. Both authors reviewed and approved the final version of the manuscript.

## Conflict of interest

The authors declare no conflicts of interest.

## Data availability statement

All raw data sequences was obtained through NCBI (see methods) and scripts for the pipeline and further analyses can be found in the following Github repository (https://github.com/PHYTOPatCAU/RLP_identification)

## Supporting information

Supplementary data 1 All SRA entries used in this study

Supplementary data 2 All predicted crude proteins for individual HMMs for variable e-values:

Supplementary data 3 Taxonomic details of Tomato and Arabidopsis

Supplementary data 4 All predicted RLPs from seven *Solanum* individuals

Supplementary data 5 Multiple sequence alignment that was used to construct the C2-C3F-HMM

Supplementary data 6 ALL 5’ coding sequence extended results

Supplementary data 7 Blastp threshold cutoff graphs Supplementary data 8 Proximity of TEs to predicted RLPs

## Notes

https://github.com/PHYTOPatCAU/RLP_identification

### Competing Interest Statement

The authors have declared no competing interest.

### Funding

- Deutsche Forschungsgemeinschaft: 516659121

## References

Andolfo G, Sanseverino W, Rombauts S, Van De Peer Y, Bradeen JM, Carputo D, Frusciante L, Ercolano MR. 2013. Overview of tomato (*Solanum lycopersicum*) candidate pathogen recognition genes reveals important *Solanum* R locus dynamics. New Phytologist 197: 223–237.

Benoit M, Jenike KM, Satterlee JW, Ramakrishnan S, Gentile I, Hendelman A, Passalacqua MJ, Suresh H, Shohat H, Robitaille GM, et al. 2025. Solanum pan-genetics reveals paralogues as contingencies in crop engineering. Nature 640: 135–145.

Bolger A, Scossa F, Bolger ME, Lanz C, Maumus F, Tohge T, Quesneville H, Alseekh S, Sørensen I, Lichtenstein G, et al. 2014. The genome of the stress-tolerant wild tomato species Solanum pennellii. Nature Genetics 46: 1034–1038.

Borowiec ML. 2016. AMAS: a fast tool for alignment manipulation and computing of summary statistics. PeerJ 4: e1660.

Camacho C, Coulouris G, Avagyan V, Ma N, Papadopoulos J, Bealer K, Madden TL. 2009. BLAST+: architecture and applications. BMC Bioinformatics 10: 421.

Capella-Gutiérrez S, Silla-Martínez JM, Gabaldón T. 2009. trimAl: a tool for automated alignment trimming in large-scale phylogenetic analyses. Bioinformatics 25: 1972–1973.

Cheng C, Krishnakumar V, Chan AP, Thibaud-Nissen F, Schobel S, Town CD. 2017. Araport11: a complete reannotation of the *Arabidopsis thaliana* reference genome. The Plant Journal 89: 789–804.

Contreras MP, Pai H, Tumtas Y, Duggan C, Yuen ELH, Cruces AV, Kourelis J, Ahn H, Lee K, Wu C, et al. 2023. Sensor NLR immune proteins activate oligomerization of their NRC helpers in response to plant pathogens. The EMBO Journal 42: EMBJ2022111519.

De La Rosa S, Schol CR, Peregrina ÁR, Winter DJ, Hilgers AM, Maeda K, Iida Y, Tarallo M, Jia R, Beenen HG, et al. 2023. Sequential breakdown of the complex Cf-9 leaf mould resistance locus in tomato by Fulvia fulva.

Dixon MS, Hatzixanthis K, Jones DA, Harrison K, Jones JDG. The Tomato Cf-5 Disease Resistance Gene and Six Homologs Show Pronounced Allelic Variation in Leucine-Rich Repeat Copy Number.

Du J, Verzaux E, Chaparro-Garcia A, Bijsterbosch G, Keizer LCP, Zhou J, Liebrand TWH, Xie C, Govers F, Robatzek S, et al. 2015. Elicitin recognition confers enhanced resistance to Phytophthora infestans in potato. Nature Plants 1: 15034.

Eddy SR. HMMER User’s Guide.

Edgar RC. 2004. MUSCLE: multiple sequence alignment with high accuracy and high throughput. Nucleic Acids Research 32: 1792–1797.

Enright AJ. 2002. An efficient algorithm for large-scale detection of protein families. Nucleic Acids Research 30: 1575–1584.

Ewels P, Magnusson M, Lundin S, Käller M. 2016. MultiQC: summarize analysis results for multiple tools and samples in a single report. Bioinformatics 32: 3047–3048.

Fritz-Laylin LK, Krishnamurthy N, Tör M, Sjölander KV, Jones JDG. 2005. Phylogenomic Analysis of the Receptor-Like Proteins of Rice and Arabidopsis. Plant Physiology 138: 611–623.

Gíslason MH, Nielsen H, Almagro Armenteros JJ, Johansen AR. 2021. Prediction of GPI-anchored proteins with pointer neural networks. Current Research in Biotechnology 3: 6–13.

Hallgren J, Tsirigos KD, Pedersen MD, Almagro Armenteros JJ, Marcatili P, Nielsen H, Krogh A, Winther O. 2022. DeepTMHMM predicts alpha and beta transmembrane proteins using deep neural networks.

Hegenauer V, Slaby P, Körner M, Bruckmüller J-A, Burggraf R, Albert I, Kaiser B, Löffelhardt B, Droste-Borel I, Sklenar J, et al. 2020. The tomato receptor CuRe1 senses a cell wall protein to identify Cuscuta as a pathogen. Nature Communications 11: 5299.

Irshad M, Canut H, Borderies G, Pont-Lezica R, Jamet E. 2008. A new picture of cell wall protein dynamics in elongating cells of Arabidopsis thaliana: Confirmed actors and newcomers. BMC Plant Biology 8: 94.

Jehle AK, Lipschis M, Albert M, Fallahzadeh-Mamaghani V, Fürst U, Mueller K, Felix G. 2013. The Receptor-Like Protein ReMAX of *Arabidopsis* Detects the Microbe-Associated Molecular Pattern eMax from *Xanthomonas*. The Plant Cell 25: 2330–2340.

Jones P, Binns D, Chang H-Y, Fraser M, Li W, McAnulla C, McWilliam H, Maslen J, Mitchell A, Nuka G, et al. 2014. InterProScan 5: genome-scale protein function classification. Bioinformatics 30: 1236–1240.

Jones JDG, Dangl JL. 2006. The plant immune system. Nature 444: 323–329.

Jones DA, Thomas CM, Hammond-Kosack KE, Balint-Kurti PJ, Jones JDG. 1994. Isolation of the Tomato *Cf-9* Gene for Resistance to *Cladosporium fulvum* by Transposon Tagging. Science 266: 789–793.

Kahlon PS, Seta SM, Zander G, Scheikl D, Hückelhoven R, Stam R. Population studies of the wild tomato species Solanum chilense reveal geographically structured major gene-mediated pathogen resistance.

Kalischuk M, Müller B, Fusaro AF, Wijekoon CP, Waterhouse PM, Prüfer D, Kawchuk L. 2022. Amplification of cell signaling and disease resistance by an immunity receptor Ve1Ve2 heterocomplex in plants. Communications Biology 5: 497.

Käll L, Krogh A, Sonnhammer ELL. 2004. A Combined Transmembrane Topology and Signal Peptide Prediction Method. Journal of Molecular Biology 338: 1027–1036.

Kang W-H, Lee J, Koo N, Kwon J-S, Park B, Kim Y-M, Yeom S-I. 2022. Universal gene co-expression network reveals receptor-like protein genes involved in broad-spectrum resistance in pepper (*Capsicum annuum* L.). Horticulture Research 9: uhab003.

Kang W-H, Yeom S-I. 2018. Genome-wide Identification, Classification, and Expression Analysis of the Receptor-Like Protein Family in Tomato. The Plant Pathology Journal 34: 435–444.

Katoh K, Standley DM. 2013. MAFFT Multiple Sequence Alignment Software Version 7: Improvements in Performance and Usability. Molecular Biology and Evolution 30: 772–780.

Kawchuk LM, Hachey J, Lynch DR, Kulcsar F, Van Rooijen G, Waterer DR, Robertson A, Kokko E, Byers R, Howard RJ, et al. 2001. Tomato *Ve* disease resistance genes encode cell surface-like receptors. Proceedings of the National Academy of Sciences 98: 6511–6515.

Kim, D., Paggi, J.M., Park, C. et al. 2019. Graph-based genome alignment and genotyping with HISAT2 and HISAT-genotype. Nat Biotechnol: 907–915.

Kim S, Park M, Yeom S-I, Kim Y-M, Lee JM, Lee H-A, Seo E, Choi J, Cheong K, Kim K-T, et al. 2014. Genome sequence of the hot pepper provides insights into the evolution of pungency in Capsicum species. Nature Genetics 46: 270–278.

Kozak M. 1978. How do eucaryotic ribosomes select initiation regions in messenger RNA? Cell 15: 1109–1123.

Krusell L, Sato N, Fukuhara I, Koch BEV, Grossmann C, Okamoto S, Oka-Kira E, Otsubo Y, Aubert G, Nakagawa T, et al. 2011. The *Clavata2* genes of pea and *Lotus japonicus* affect autoregulation of nodulation. The Plant Journal 65: 861–871.

Leister D. 2004. Tandem and segmental gene duplication and recombination in the evolution of plant disease resistance genes. Trends in Genetics 20: 116–122.

Li H, Handsaker B, Wysoker A, Fennell T, Ruan J, Homer N, Marth G, Abecasis G, Durbin R, 1000 Genome Project Data Processing Subgroup, et al. 2009. The Sequence Alignment/Map format and SAMtools. Bioinformatics 25: 2078–2079.

Li H-L, Wu L, Dong Z, Jiang Y, Jiang S, Xing H, Li Q, Liu G, Tian S, Wu Z, et al. 2021. Haplotype-resolved genome of diploid ginger (*Zingiber officinale*) and its unique gingerol biosynthetic pathway. Horticulture Research 8: 189.

Macho AP, Zipfel C. 2014. Plant PRRs and the Activation of Innate Immune Signaling. Molecular Cell 54: 263–272.

Manni M, Berkeley MR, Seppey M, Zdobnov EM. 2021. BUSCO: Assessing Genomic Data Quality and Beyond. Current Protocols 1: e323.

Martin RA, Tate AT. 2024. Pleiotropy alleviates the fitness costs associated with resource allocation trade-offs in immune signalling networks. Proceedings of the Royal Society B: Biological Sciences 291: 20240446.

Masi A, Trentin AR, Arrigoni G. 2016. Leaf apoplastic proteome composition in UV-B treated Arabidopsis thaliana mutants impaired in extracellular glutathione degradation. Data in Brief 6: 368–377.

Michelmore RW, Meyers BC. 1998. Clusters of Resistance Genes in Plants Evolve by Divergent Selection and a Birth-and-Death Process. Genome Research 8: 1113–1130.

Minh BQ, Schmidt HA, Chernomor O, Schrempf D, Woodhams MD, Von Haeseler A, Lanfear R, Teeling E. 2020. IQ-TREE 2: New Models and Efficient Methods for Phylogenetic Inference in the Genomic Era (E Teeling, Ed.). Molecular Biology and Evolution 37: 1530–1534.

Mirarab S, Reaz R, Bayzid MdS, Zimmermann T, Swenson MS, Warnow T. 2014. ASTRAL: genome-scale coalescent-based species tree estimation. Bioinformatics 30: i541–i548.

Muir CD, Pease JB, Moyle LC. 2014. Quantitative Genetic Analysis Indicates Natural Selection on Leaf Phenotypes Across Wild Tomato Species (*Solanum* sect. Lycopersicon; Solanaceae). Genetics 198: 1629–1643.

Ngou BPM, Ding P, Jones JDG. 2022a. Thirty years of resistance: Zig-zag through the plant immune system. The Plant Cell 34: 1447–1478.

Ngou BPM, Heal R, Wyler M, Schmid MW, Jones JDG. 2022b. Concerted expansion and contraction of immune receptor gene repertoires in plant genomes. Nature Plants 8: 1146–1152.

Ngou BPM, Wyler M, Schmid MW, Kadota Y, Shirasu K. 2024. Evolutionary trajectory of pattern recognition receptors in plants. Nature Communications 15: 308.

Nielsen H, Kihara D. 2017. Predicting Secretory Proteins with SignalP. In: Kihara D, ed. Protein Function Prediction. New York, NY: Springer New York, 59–73.

Ou S, Su W, Liao Y, Chougule K, Agda JRA, Hellinga AJ, Lugo CSB, Elliott TA, Ware D, Peterson T, et al. 2019. Benchmarking transposable element annotation methods for creation of a streamlined, comprehensive pipeline. Genome Biology 20: 275.

Parniske M, Jones JDG. 1999. Recombination between diverged clusters of the tomato *Cf-9* plant disease resistance gene family. Proceedings of the National Academy of Sciences 96: 5850–5855.

Pease JB, Haak DC, Hahn MW, Moyle LC, Penny D. 2016. Phylogenomics Reveals Three Sources of Adaptive Variation during a Rapid Radiation (D Penny, Ed.). PLOS Biology 14: e1002379.

Pei L, Wang B, Ye J, Hu X, Fu L, Li K, Ni Z, Wang Z, Wei Y, Shi L, et al. 2021. Genome and transcriptome of Papaver somniferum Chinese landrace CHM indicates that massive genome expansion contributes to high benzylisoquinoline alkaloid biosynthesis. Horticulture Research 8: 5.

Penczykowski RM, Stam R, Halliday FW. Bridging plant pathology and disease ecology: emerging insights into wild plant–pathogen systems.

Peterson KM, Rychel AL, Torii KU. 2010. Out of the Mouths of Plants: The Molecular Basis of the Evolution and Diversity of Stomatal Development. The Plant Cell 22: 296–306.

Powell AF, Feder A, Li J, Schmidt MH -W., Courtney L, Alseekh S, Jobson EM, Vogel A, Xu Y, Lyon D, et al. 2022. A *Solanum lycopersicoides* reference genome facilitates insights into tomato specialized metabolism and immunity. The Plant Journal 110: 1791–1810.

Price MN, Dehal PS, Arkin AP. 2009. FastTree: Computing Large Minimum Evolution Trees with Profiles instead of a Distance Matrix. Molecular Biology and Evolution 26: 1641–1650.

Quinlan AR, Hall IM. 2010. BEDTools: a flexible suite of utilities for comparing genomic features. Bioinformatics 26: 841–842.

Rameneni JJ, Lee Y, Dhandapani V, Yu X, Choi SR, Oh M-H, Lim YP, Raman H. 2015. Genomic and Post-Translational Modification Analysis of Leucine-Rich-Repeat Receptor-Like Kinases in Brassica rapa (H Raman, Ed.). PLOS ONE 10: e0142255.

Robinson JT. 2011. Integrative genomics viewer. correspondence 29.

Ron M, Avni A. 2004. The Receptor for the Fungal Elicitor Ethylene-Inducing Xylanase Is a Member of a Resistance-Like Gene Family in Tomato. The Plant Cell 16: 1604–1615.

Seifbarghi S, Borhan MH, Wei Y, Ma L, Coutu C, Bekkaoui D, Hegedus DD. 2020. Receptor-Like Kinases BAK1 and SOBIR1 Are Required for Necrotizing Activity of a Novel Group of Sclerotinia sclerotiorum Necrosis-Inducing Effectors. Frontiers in Plant Science 11: 1021.

Silva-Arias GA, Gagnon E, Hembrom S, Fastner A, Khan MR, Stam R, Tellier A. 2025. Patterns of presence–absence variation of NLRS across populations of *Solanum chilense* are clade-dependent and mainly shaped by past demographic history. New Phytologist 245: 1718–1732.

Snoeck S, Garcia AGk, Steinbrenner AD. 2023. Plant Receptor-like proteins (RLPs): Structural features enabling versatile immune recognition. Physiological and Molecular Plant Pathology 125: 102004.

Sonnhammer ELL, Koonin EV. 2002. Orthology, paralogy and proposed classification for paralog subtypes. Trends in Genetics 18: 619–620.

Stam R, Nosenko T, Hörger AC, Stephan W, Seidel M, Kuhn JMM, Haberer G, Tellier A. 2019. The *de Novo* Reference Genome and Transcriptome Assemblies of the Wild Tomato Species *Solanum chilense* Highlights Birth and Death of NLR Genes Between Tomato Species. G3 Genes|Genomes|Genetics 9: 3933–3941.

Stamatakis A. 2014. RAxML version 8: a tool for phylogenetic analysis and post-analysis of large phylogenies. Bioinformatics 30: 1312–1313.

Steidele CE, Stam R. 2021. Multi-omics approach highlights differences between RLP classes in Arabidopsis thaliana. BMC Genomics 22: 557.

Steuernagel B, Witek K, Krattinger SG, Ramirez-Gonzalez RH, Schoonbeek H, Yu G, Baggs E, Witek AI, Yadav I, Krasileva KV, et al. 2020. The NLR-Annotator Tool Enables Annotation of the Intracellular Immune Receptor Repertoire. Plant Physiology 183: 468–482.

Sutherland CA, Prigozhin DM, Monroe JG, Krasileva KV. 2024. High allelic diversity in Arabidopsis NLRs is associated with distinct genomic features. EMBO Reports 25: 2306–2322.

The UniProt Consortium, Bateman A, Martin M-J, Orchard S, Magrane M, Adesina A, Ahmad S, Bowler-Barnett EH, Bye-A-Jee H, Carpentier D, et al. 2025. UniProt: the Universal Protein Knowledgebase in 2025. Nucleic Acids Research 53: D609–D617.

Thomas CM, Hatzixanthis K, Laboratory S. Characterization of the Tomato Cf-4 Gene for Resistance to Cladosporium fulvum ldentifies Sequences That Determine RecognitionalSpecificity in Cf-4 and Cf-9.

Torres Ascurra YC, Zhang L, Toghani A, Hua C, Rangegowda NJ, Posbeyikian A, Pai H, Lin X, Wolters PJ, Wouters D, et al. 2023. Functional diversification of a wild potato immune receptor at its center of origin. Science 381: 891–897.

Van De Weyer A-L, Monteiro F, Furzer OJ, Nishimura MT, Cevik V, Witek K, Jones JDG, Dangl JL, Weigel D, Bemm F. 2019. A Species-Wide Inventory of NLR Genes and Alleles in Arabidopsis thaliana. Cell 178: 1260–1272.e14.

Van Der Burgh AM, Postma J, Robatzek S, Joosten MHAJ. 2019. Kinase activity of SOBIR1 and BAK1 is required for immune signalling. Molecular Plant Pathology 20: 410–422.

Van Der Hoorn RAL, Kruijt M, Roth R, Brandwagt BF, Joosten MHAJ, De Wit PJGM. 2001. Intragenic recombination generated two distinct *Cf* genes that mediate AVR9 recognition in the natural population of *Lycopersicon pimpinellifolium*. Proceedings of the National Academy of Sciences 98: 10493–10498.

Wang G, Ellendorff U, Kemp B, Mansfield JW, Forsyth A, Mitchell K, Bastas K, Liu C-M, Woods-Tör A, Zipfel C, et al. 2008. A Genome-Wide Functional Investigation into the Roles of Receptor-Like Proteins in Arabidopsis. Plant Physiology 147: 503–517.

Wu Y, Xun Q, Guo Y, Zhang J, Cheng K, Shi T, He K, Hou S, Gou X, Li J. 2016. Genome-Wide Expression Pattern Analyses of the Arabidopsis Leucine-Rich Repeat Receptor-Like Kinases. Molecular Plant 9: 289–300.

Wulff BBH, Thomas CM, Smoker M, Grant M, Jones JDG. Domain Swapping and Gene Shuffling Identify Sequences Required for Induction of an Avr-Dependent Hypersensitive Response by the Tomato Cf-4 and Cf-9 Proteins.

Yang H, Bayer PE, Tirnaz S, Edwards D, Batley J. 2020. Genome-Wide Identification and Evolution of Receptor-Like Kinases (RLKs) and Receptor like Proteins (RLPs) in Brassica juncea. Biology 10: 17.

Krueger F, James F, Ewels P, Afyounian E, Schuster-Boeckler B. 2023. TrimGalore: v0.6.10.

