## Supplementary tables and figures for "PlantLRR-PRR: A Reproducible Annotation Pipeline Reveals Contrasting Evolution of Developmental and Defense Receptor-Like Proteins in Tomato"

#### Supplementary Table 1: Proteome data, primary gene counts, and BUSCO values of all the plant genomes used in this study.

| *Solanum spp* | Source | Total protein count | proteins in primary gene models | BUSCO completeness (percentage) | Weblink to download |
|---|---|---|---|---|---|
| *S. pennellii_0716* | Solgenomics.net | 47679 | 44965 | 99.3 | https://solgenomics.net/ftp/genomes/Solanum_pennellii/ |
| *S. lycopersicum (ITGA.2.4)* | Solgenomics.net | 34725 | 34725 | 99.4 | https://solgenomics.net/ftp/genomes/Solanum_lycopersicum/annotation/ITAG2.4_release/ |
| *S. pimpinellifolium _1589* | Solgenomics.net | 41449 | 41449 | 99.5 | https://doi.org/10.6084/m9.figshare.24605586 |
| *S. pimpinellifolium _BGV06775* | Solgenomics.net | 35111 | 34465 | 99.4 | https://solgenomics.net/ftp/genomes/Solanum_pimpinellifolium/BGV006775/ |
| *S. pimpinellifolium _PAS014479* | Solgenomics.net | 35083 | 35083 | 99.5 | https://solgenomics.net/ftp/genomes/TGG/genome/PAS014479.fasta.gz |
| *S. pimpinellifolium _PI365967* | Solgenomics.net | 35360 | 35360 | 99.5 | https://solgenomics.net/ftp/genomes/TGG/genome/PP.fasta.gz |
| *S. lycopersicoides* | Solgenomics | 37938 | 34207 | 99.3 | https://solgenomics.net/ftp/genomes/Solanum_lycopersicoides/ |
| *S. chilense* | Not published yet | 86465 | 86465 | 99.3 | Not published yet |
| *A. thaliana* | Phytozome13 | 27654 | 27654 | 99.9 | https://phytozome-next.jgi.doe.gov/info/Athaliana_Araport11 |
| *A. halleri* | Phytozome13 | 28722 | 28722 | 99.6 | https://phytozome-next.jgi.doe.gov/info/Ahalleri_v2_1_0 |
| *S. tuberosum* | Phytozome13 | 32917 | 32917 | 99.6 | https://phytozome-next.jgi.doe.gov/info/Stuberosum_v6_1 |
| *P. vulgaris* | Phytozome13 | 27385 | 27385 | 99.2 | https://phytozome-next.jgi.doe.gov/info/Pvulgaris_v2_1 |

#### Supplementary Table 2: List of Pfam IDs for the respective LRR, Kinase, and NLR domains.

| **LRR Pfam IDs (Pfam33.0)** | | | | **NLR Pfam IDs** | | **Pkinase Pfam IDs** | |
|---|---|---|---|---|---|---|---|
| Domains | IDs | Domains | IDs | Domains | IDs | Domains | IDs |
| LRR-1 | PF00560.34 | LRR-11 | PF18831.2 | ABC | PF03109.17 | PK_Tyr_Ser_Thr | PF07714.18 |
| LRR-2 | PF07723.14 | LRR-12 | PF18837.2 | TIR | PF01582.21 | Pkinase | PF00069.26 |
| LRR-3 | PF07725.13 | LRRNT | PF01462.19 | TIR-2 | PF13676.7 | Pkinase-C | PF00433.25 |
| LRR-4 | PF12799.8 | LRRCT | PF01463.25 | TIR-3 | PF18567.2 | | |
| LRR-5 | PF13306.7 | | | NB-ARC | PF00931.23 | | |
| LRRNT-2 | PF08263.13 | | | NB-LRR | PF12061.9 | | |
| LRR-6 | PF13516.7 | | | AAA-2 | PF13401.7 | | |
| LRR-8 | PF13855.7 | | | C-JID | PF20160.3 | | |
| LRR-9 | PF14580.7 | | | RPW8 | PF05659.12 | | |
| LRR-10 | PF18805.2 | | | RxN | PF18052.2 | | |

#### Supplementary Table 3: List of annotated RLP/RLKs in the previous studies.

| **Plant species** | **Previous studies** | | **Best source** |
|---|---|---|---|
| | **RLKs** | **RLPs** | |
| ***A. thaliana*** | 223 | 57 | (Shiu & Bleecker, n.d.; Wang et al., 2008)[12], [59] |
| ***A. thaliana*** | 213 | 54 | (Ngou et al., 2024) [5] |
| ***A. halleri*** | 167 | 46 | (Ngou et al., 2024) [5] |
| ***S. lycopersicum*** | 223 | 48 | (Ngou et al., 2024) [5] |
| ***S. tuberosum*** | 242 | 122 | (Ngou et al., 2024) [5] |
| ***P. vulgaris*** | 238 | 91 | (Ngou et al., 2024) [5] |

#### Supplementary Table 4: List of extracted RLPs and RLKs from all the plant genotypes used in this study

| **Plant species** | **RLKs** | **RLPs** |
|---|---|---|
| *A. thaliana* | 224 | 60 |
| *A. halleri (v2.1.0)* | 236 | 45 |
| *S.tuberosum* | 301 | 145 |
| *P. vulgaris* | 266 | 95 |
| *S.pennellii (0716)* | 232 | 60 |
| *S.lycopersicum (ITGA)* | 223 | 67 |
| *S.pimpinellifolium (1589)* | 235 | 60 |
| *S.pimpinellifolium (2093)* | 197 | 51 |
| *S.pimpinellifolium (BVG006775)* | 216 | 59 |
| *S.pimpinellifolium (PAS014479)* | 225 | 67 |
| *S.pimpinellifolium (PI365967)* | 223 | 67 |
| *S.chilense* | 239 | 93 |
| *S.lycopersicoides* | 258 | 94 |

**Supplementary Table 5: Statistical results of testing the significance of TE distances with defence and developmental RLPs**

| Component | Effect | Estimate (interpretation) | p-value | Remarks |
|---|---|---|---|---|
| sModel fit | Zero-inflated Tweedie | 3183.45 | | Better fit |
| | Non-zero-inflated Tweedie | 3203.18 | | |
| | ΔAIC | 19.73 | | Zero-inflated model is strongly preferred (>10.0) |
| Conditional (Tweedie) component | Multiplicative effect (Development vs Defence) | 0.936 | 0.589 | No significant difference in non-zero TE distance between defence and developmental gene types. |
| Zero-inflation component | Odds ratio (development vs defence) | 0.324 | $<1x10^{-7}$ | Development genes have ~3x lower odds of being zero distance/fused with TEs |
| | Estimated zero-distance probability (defence genes) | ~0.73 | | 73% of defence genes are likely fused with TEs (zero-distance) |
| | Estimated zero-distance probability (development genes) | ~0.47 | | 47% of the development genes likely fused with TEs (Zero-distance) |
| Chi-Square test | Independent or dependent test | $X^2$ = 24.27 | $5.83x10^{-7}$ | There is a dependency of TE proximity to the gene clusters |
| Fisher's exact test | Odds ratio | OR = 0.33 | $9.19x10^{-7}$ | Developmental genes are ~3x less likely to have overlapped with TEs compared to defence-related genes. |

### Supplementary figures

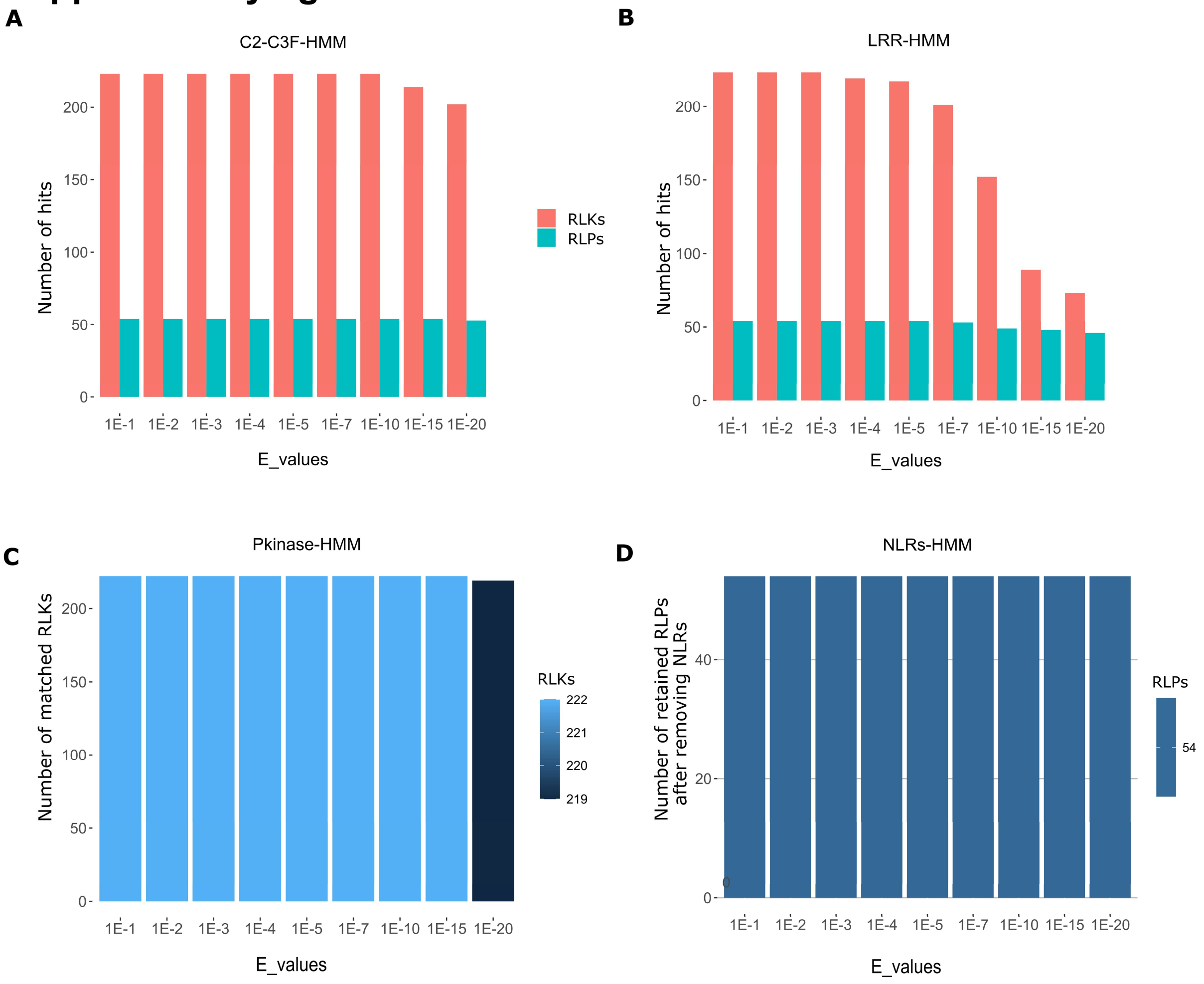


***Supplementary figure-1: Individual HMMs extracted RLPs and RLKs compared against known 57/223 RLPs/RLKs of A. thaliana under variable e-values (1E-1 to 1E-20).***

*Number of matched genome-wide extracted crude RLPs and RLKs from A. thaliana primary proteome against putatively known 57/223 RLPs and RLKs under A) C2-C3F-HMM, B) LRR-HMM, C) Pkinase-HMM models. D) No. of retained RLPs matched against 57 known RLPs after removing potential NLRs.*

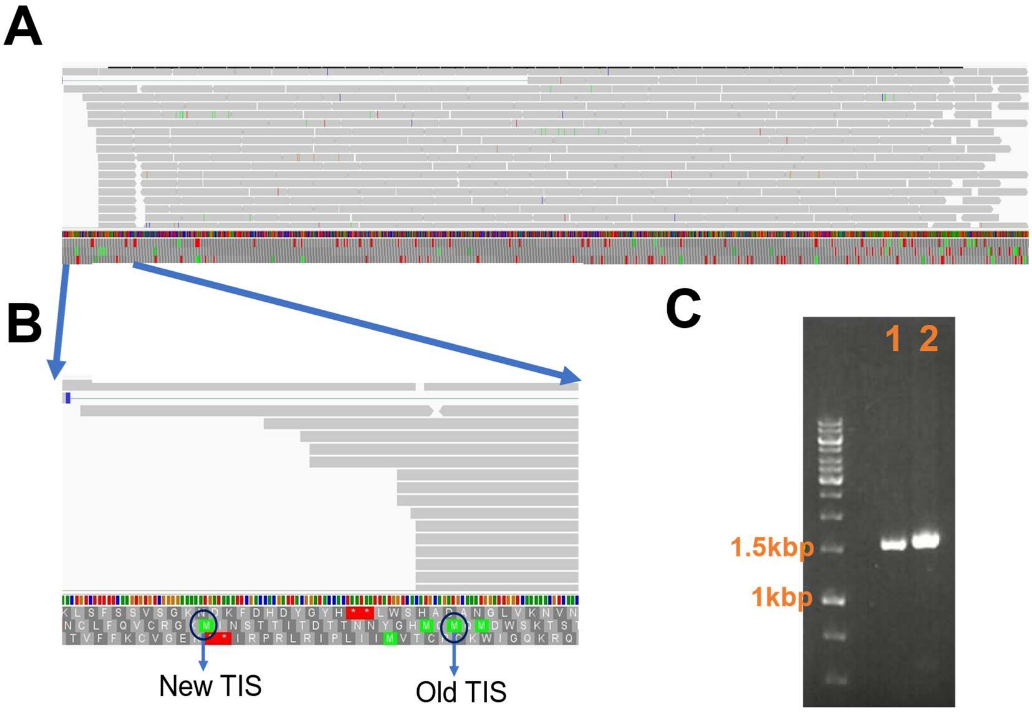


***Supplementary figure 2: Visualization and validation of transcriptomic signature in the extended region of Arabidopsis's AT1G33590.2 RLP.***

*A) RNA-seq alignment of A. thalina Col-0 against extended AT1G33590.2 mRNA. a) Raw coverage track, b) RNA-reads aligned track, c) reference consensus mRNA track, d) translation track for the reference sequence. B) Snapshot of the RNA-reads aligned to the extended region of AT1G33590.2 mRNA. Old TIS is the old translation initiation start. New TIS is the new TIS in the extended region. Red and green boxes were the stop and methionine (start) codons, respectively.C) Successful amplification of both old, "1" (1545bp), and new-extended, "2" (1599bp), coding sequences of the AT1G33590.3 gene.*

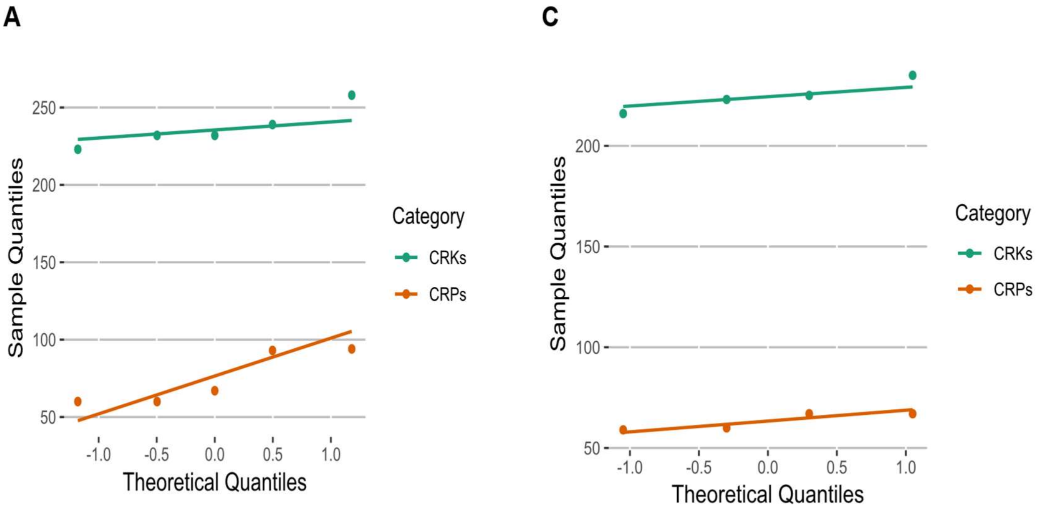


B

| Category | W | P-value | CV |
|---|---|---|---|
| CRKs | 0.8057 | 0.0901 | 0.0422 |
| CRPs | 0.7978 | 0.0777 | 0.2275 |

D

| Category | W | P-value | CV |
|---|---|---|---|
| CRKs | 0.9705 | 0.8449 | 0.0349 |
| CRPs | 0.7821 | 0.0739 | 0.0687 |

***Supplementary figure 3: Normality and the coefficient of variation of the extracted CRKs and CRPs.***

*A-B) Normality was checked between 5 Solanum spp, Q-Q plot displaying the distribution of CNV of CRKs and CRPs, and Shapiro wilk test result, "W", proved to be closure to 1 with P-value >0.05 for both CRPs and CRKs.*

*C-D) Normality was checked between 4 Solanum pimpinellifolium, Q-Q plot displaying the distribution of CNV of CRKs and CRPs, and Shapiro wilk test result, "W", proved to be closure to 1 with P-value >0.05 for both CRPs and CRKs.*

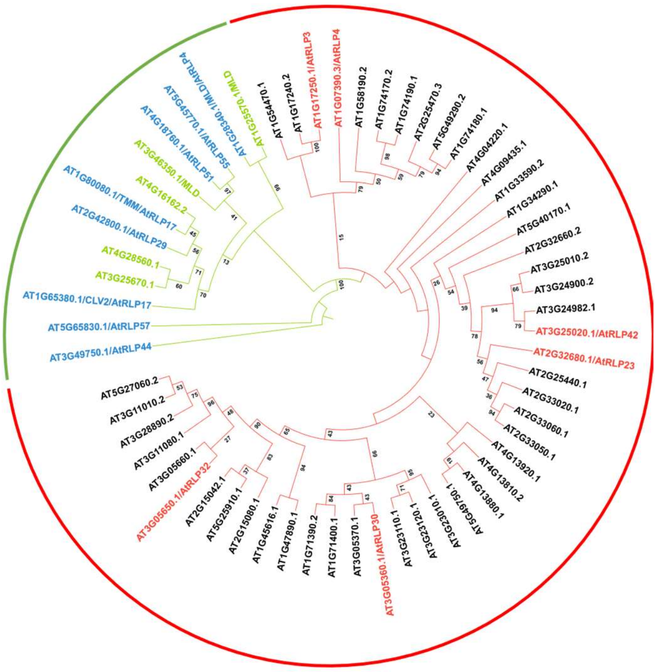


***Supplementary Figure 4: Phylogenetic reconstruction of RLPs in Arabidopsis thaliana.***

*Phylogenetic tree was constructed by targeting C2-C3F domain from 60 predicted RLP from Arabidopsis thaliana. The tree was generated using RAxML with 1000 bootstrap replicates. Developmental and defense-related RLPs, following Steidele et al. (2021), segregate into two well-supported superclades, highlighted in green (developmental) and red (defense). The tree was rooted on the developmental RLP RLP57. Bootstrap support for the split between the two superclades was 100%. Known putative developmental RLPs included RLP4, 10/CLV2, 17/TMM, 29, 44, 51, 55, and 57; known defense-related RLPs included RLP1, 3, 23, 30, 32, and 42..*

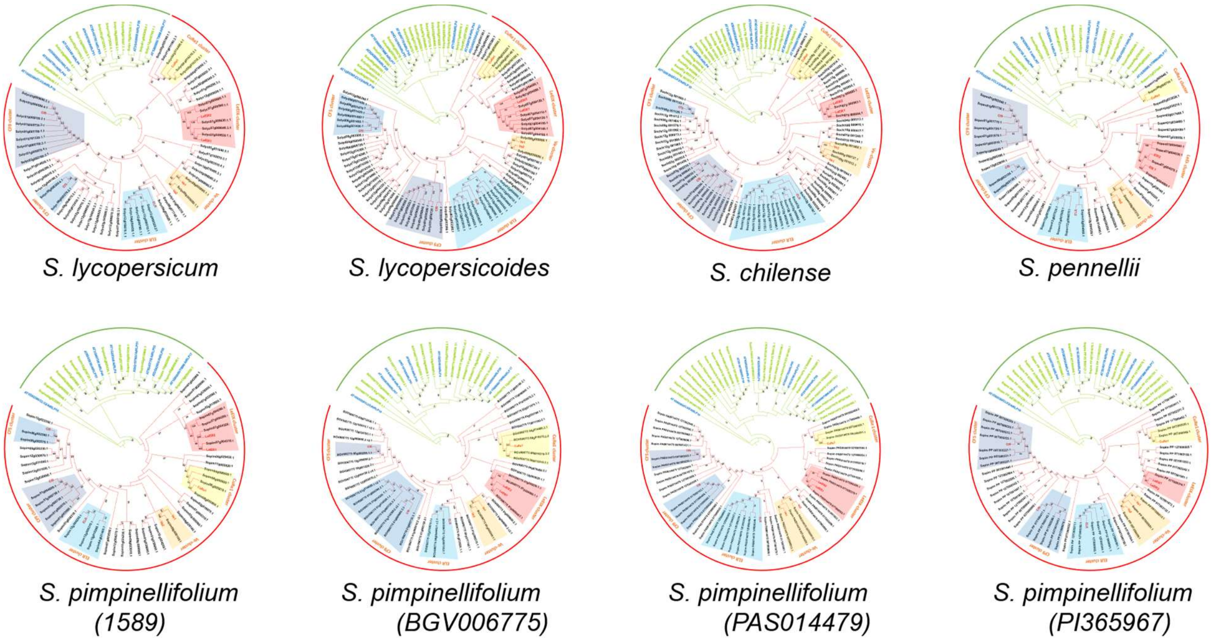


***Supplementary fig-5: Individual phylogenetic trees of predicted CRPs from all 8 Solanum genotypes.***

*Individaul phylogenetic tree were constructed by targeting prominent C2C3F domain from all the predicted CRPs from 8 Solanum genotypes. Sixteen known CRPs with established functionalities were included as referece: developmental CRPs (blue labeled) and defense-related CRPs (red labeled). RLP17/TMM, 51, 55, 29, 4, 10/CLV2, 44, and 57 were the eight known development-related CRPs, while, Cf9, Cf5, ELR, Ve1, Ve2, EiX1, EiX2, and CuRe1 were the eight known defense-related CRPs. The tree was inferred with RAxML using 1000 bootstraps. Two well-suported superclades, similar to Supplementary figure-1, (100% bootstrap strength) corresponding to developmental and defense-related CRPs were indicated by green and red acrs, respectively.*

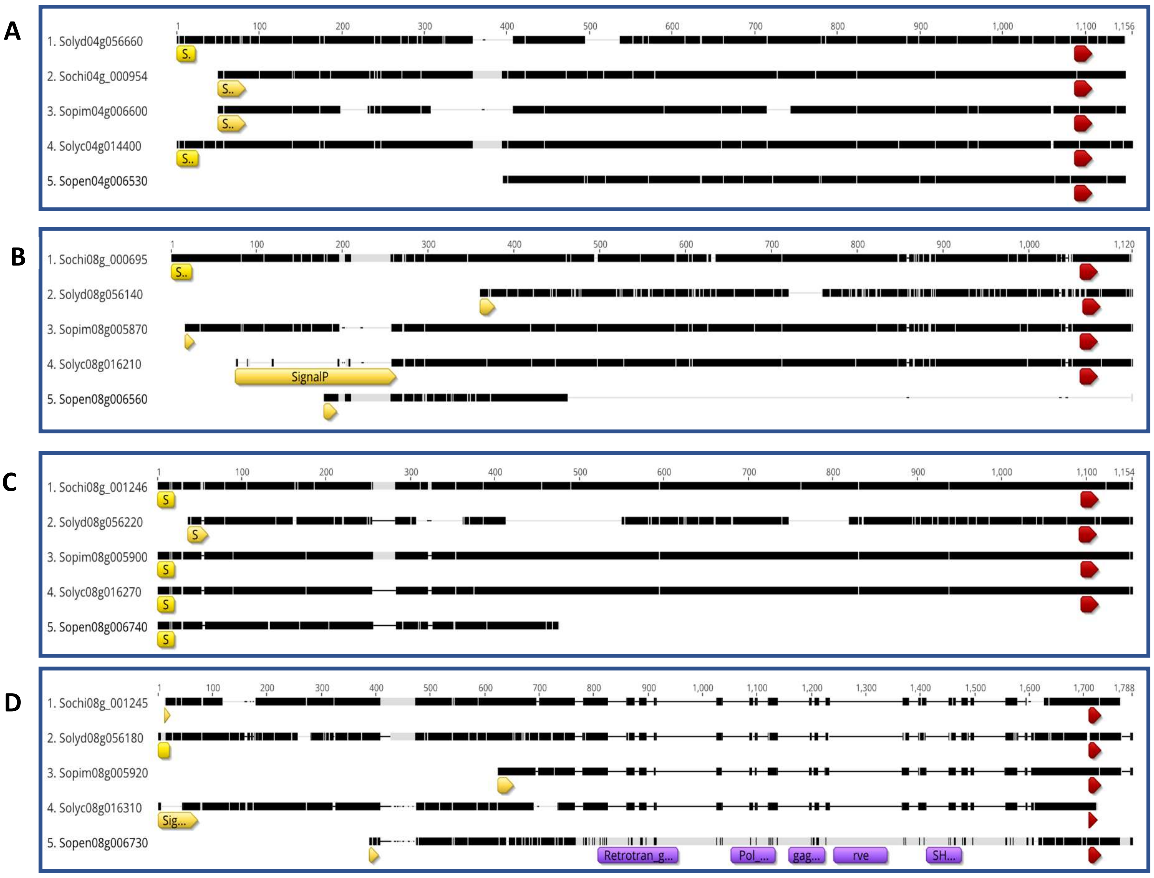


***Supplementary fig-6: Multiple sequence alignment of CuRe1 orthogroup cluster between five Solanum species.***

*Cure1 cluster in each Solanum species comprised of 4 paralogs combined to form CuRe1 orthogroup.*
*A) MSA of Solyc04g01440 orthologs between 5 Solanum species.*
*B) MSA of Solyc08g016210 orthologs between 5 Solanum species.*
*C) MSA of Solyc08g016270 orthologs between 5 Solanum species.*
*D) MSA of Solyc08g016310 orthologs between 5 Solanum species.*

**Yellow and red blocks represent annotated putative signal-peptide and transmembrane domain, respectively.*
**Purple blocks represent the annotated transposable element; Retrotransposon and its elements, in Solanum pennellii CRP, Sopen08g006730.*

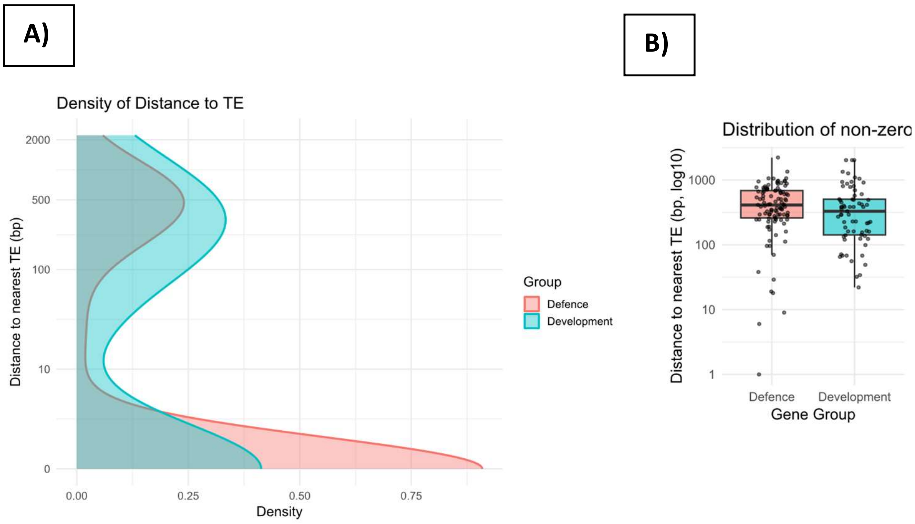


***Supplementary Figure 7: Transposable elements were in proximity to defense-related CRPs compared to developmental-related CRPs***

*A) Raw density distribution of CRPs type with respect to their nearest TE.*
*B) Mean distance of CRPs types accounting only for non-zero TE distances.*

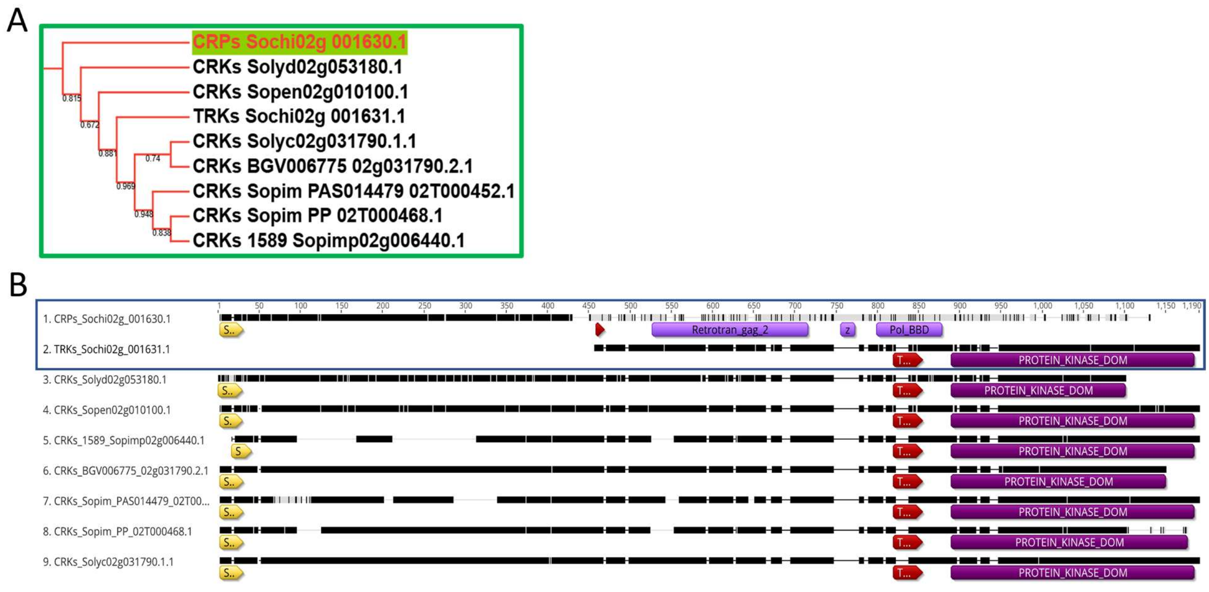


***Supplementary Figure 8: Insertion of a transposable element divided the complete RLK (CRK) into potential complete RLp (CRPs) and truncated RLK (TRKs) in S. chilense.***
*A) Local branch of a phylogenetic tree extracted with Orthologs belongs to CRKs loci, Solyc02g031790.1, shared among all 8 Solanum genotypes.*
*B) Multiple sequence alignment and annotation of orthologs belong to the CRKs loci, Solyc02g031790.1. Yellow, red, dark-pink, and purple boxes displaying SP, TMD, kinase domain, and elements of retrotransposons, respectively*
