## Supplementary figures and images for "PlantLRR-PRR: A Reproducible Annotation Pipeline Reveals Contrasting Evolution of Developmental and Defense Receptor-Like Proteins in Tomato"

### Supplementary data 5

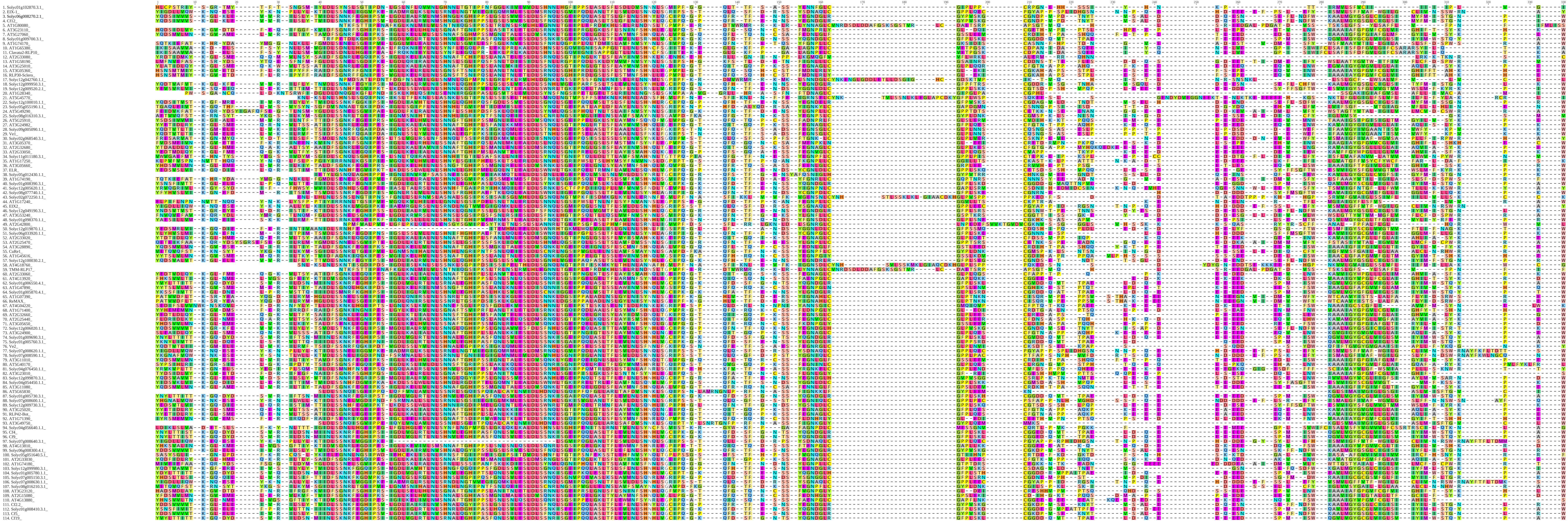
