## Supplementary data 7 for "PlantLRR-PRR: A Reproducible Annotation Pipeline Reveals Contrasting Evolution of Developmental and Defense Receptor-Like Proteins in Tomato"

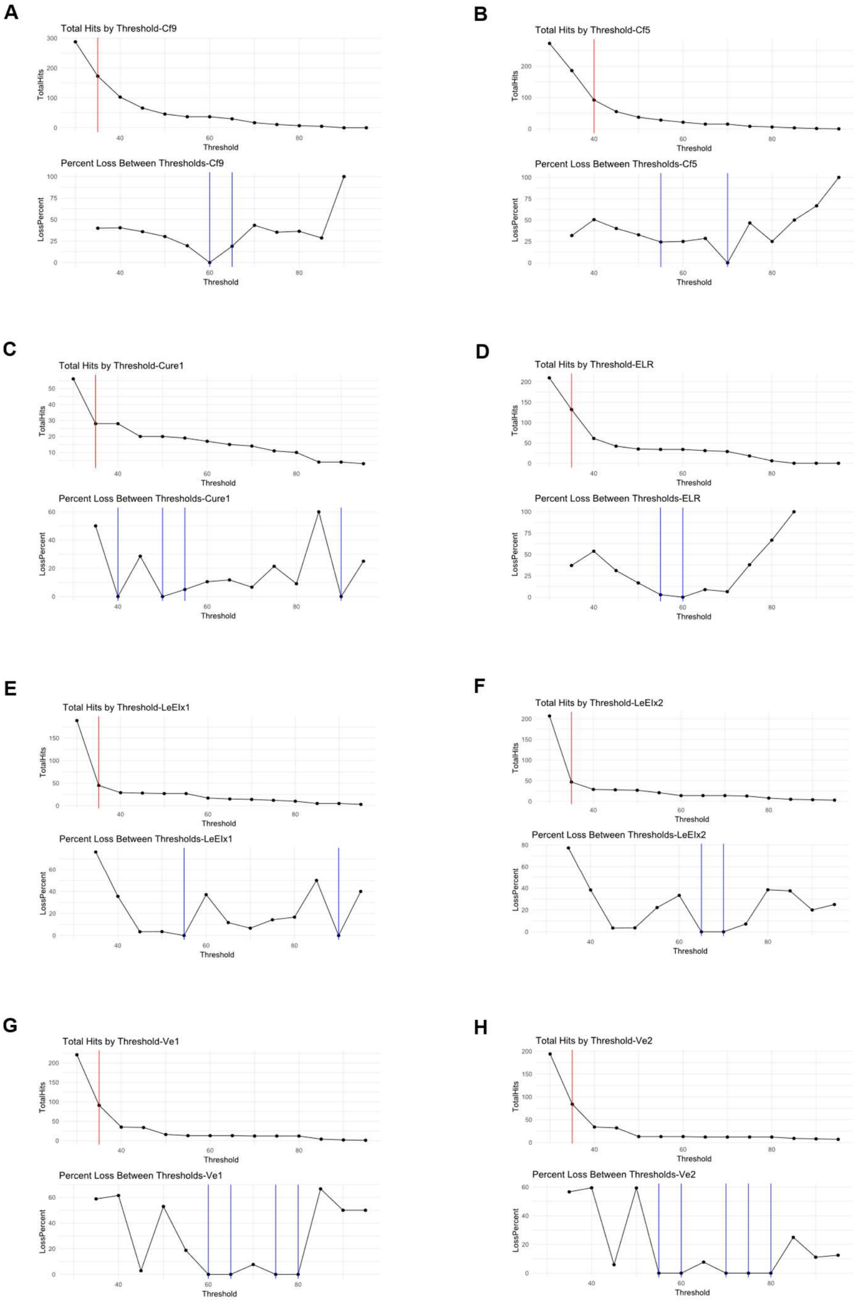


Figure 1: Sequence identity cutoff graphs for defence-related RLP orthogrouping. A-H) Graphs showing total hits for each sequence identity threshold between 30% till 95% with 5% increments and percent loss between two consecutive thresholds for individual query sequences of Cf9, Cf5, CuRe1, ELR, LeEIX1, LeEIX2, Ve1, and Ve2, respectively. Red line, the elbow point for finding the sequence identity threshold that keeps all related hits, after which we calculate the percent loss between consecutive thresholds. Blue lines showing the first and the second lowest loss recorded for the differences between two consecutive percentage identity thresholds.

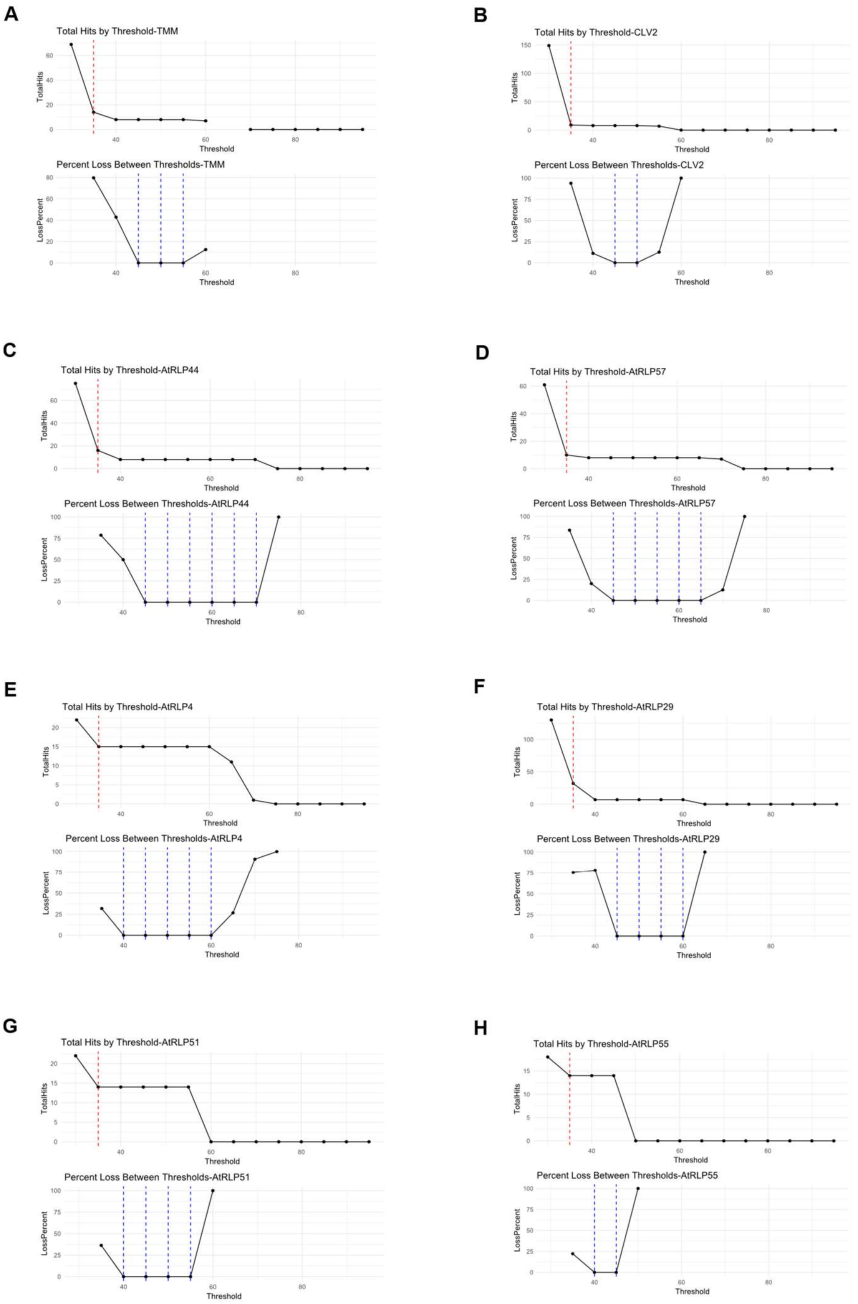


Figure 2: Sequence identity cutoff graphs for developmental-related RLP orthogrouping. A-H) Graphs showing total hits for each sequence identity threshold between 30% till 95% with 5% increments and percent loss between two consecutive thresholds for individual query sequences of TMMs, CLV2, RLP44, RLP57, RLP4, RLP29, RLP51, and RLP55, respectively. Red line, the elbow point for finding the sequence identity threshold that keeps all related hits, after which we calculate the percent loss between consecutive thresholds. Blue lines showing the first and the second lowest loss recorded for the differences between two consecutive percentage identity thresholds.
